# Minimum epistasis interpolation for sequence-function relationships

**DOI:** 10.1101/657841

**Authors:** Juannan Zhou, David M. McCandlish

## Abstract

Massively parallel phenotyping assays have provided unprecedented insight into how multiple mutations combine to determine biological function. While these assays can measure phenotypes for thousands to millions of genotypes in a single experiment, in practice these measurements are not exhaustive, so that there is a need for techniques to impute values for genotypes whose phenotypes are not directly assayed. Here we present a method based on the idea of inferring the least epistatic possible sequence-function relationship compatible with the data. In particular, we infer the reconstruction in which mutational effects change as little as possible across adjacent genetic backgrounds. Although this method is highly conservative and has no tunable parameters, it also makes no assumptions about the form that genetic interactions take, resulting in predictions that can behave in a very complicated manner where the data require it but which are nearly additive where data is sparse or absent. We apply this method to analyze a fitness landscape for protein G, showing that our technique can provide a substantially less epistatic fit to the landscape than standard methods with little loss in predictive power. Moreover, our analysis reveals that the complex structure of epistasis observed in this dataset can be well-understood in terms of a simple qualitative model consisting of three fitness peaks where the landscape is locally additive in the vicinity of each peak.

## Introduction

Recent advances in quantification via next-generation sequencing have allowed the proliferation of high-throughput combinatorial mutagenesis assays that measure molecular function for tens of thousands up to millions of sequences simultaneously [1]. These assays have been applied to many different classes of functional elements, including protein coding sequences [2–13], RNAs [14–18], and regulatory or splicing elements [19–22]. However, in practice, due to both the vastness of sequence space and the limitations of techniques for library preparation, such experiments typically result in missing measurements for a subset of possible genotypes.

Making accurate phenotypic predictions for these missing sequences is a difficult problem because the effect of any given mutation often depends on which other mutations are already present in the sequence, a phenomenon known as epistasis [23–25]. In the special case where such interactions are limited to occurring between pairs of sites, the prediction problem can be solved using regularized regression [26] a technique that has sometimes performed quite well [27, 28]. However, there is now abundant evidence that adding pair-wise interaction terms to an otherwise additive model is not sufficient to capture the complex interdependencies between mutations observed in empirical data [10, 24, 29–39].

In principle, these “higher-order” interactions can be captured by adding interactions between three or more sites to standard regression models, but this leads to problems in interpretability and overfitting because the number of such terms grows rapidly with increasing interaction order [26]. Another strategy has been to assume that the observed phenotype is a simple non-linear function of some underlying non-epistatic trait [32, 40], a pattern of epistasis known as univariate [8, 24], non-specific [31] or global [40, 41] epistasis, which appears to be well suited-primarily to sequence-function relationships that are essentially noised versions of single-peaked landscapes. Finally, a variety of machine-learning techniques [8, 12, 42–45] have been employed that can fit more complex forms of epistasis than global epistasis or pairwise interaction models. However, these require substantial tuning and the resulting models exhibit behavior that is difficult to interpret.

Here we present a new method for fitting sequence-function relationships that includes epistatic interactions of all orders but whose predictions are nonetheless conservative, which has no tunable parameters, and which is simple enough to provide formal mathematical guarantees on its behavior. The main idea is to assign the missing phenotypic values in such a way that the effects of mutations are as consistent across mutationally adjacent genetic backgrounds as possible. We achieve this by minimizing the expected squared epistatic coefficient for random pairs of mutations over all possible genetic backgrounds, a minimization problem that comes down to solving a single set of coupled linear equations. The end result is a model that can provide a complicated fit where data is abundant, but which approaches additivity in regions of sequence space where data is sparse or absent.

In what follows, we first describe our modeling technique and its mathematical properties. We then compare our method with regression models in terms of predictive power and behavior, using a simulated dataset under a simple biophysical model for transcriptional regulation [46], and data from an experimental deep mutational scanning study of the protein domain GB1 [34]. Finally, we combine our imputation procedure and a previously proposed visualization technique [47] to identify the major qualitative features of the sequence-function relationship for GB1 that determine and explain the complex patterns of genetic interaction previously observed in this dataset.

## Minimum epistasis interpolation

Given fitness observations on a subset of genotypes, our goal is to assign fitness values to all unobserved genotypes in such a manner that mutational effects change as little as possible between mutationally adjacent genetic backgrounds. To understand our solution to this problem, it is helpful to think about the simplest possible case where sequence space consists of two bi-allelic loci and hence four possible genotypes (WT, ab; single mutants aB, Ab; and the double mutant, AB). We assume we have observed phenotypes for the WT and both single mutants, and wish to predict the phenotype of the double mutant.

For this simple case, we can measure the change in the effect of a mutation at the first locus (a → A) between the two backgrounds at the second locus using the traditional epistatic coefficient [48]

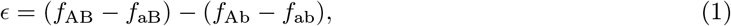

which is just the difference in the effect of an a → A mutation between the *b* and *B* backgrounds, Figure 1A. Now, the sign of *ϵ* depends on which sequence we have assigned to be WT, so if we want a reference-free measure of how much mutational effects change with genetic background we can instead use the squared quantity *ϵ*^2^, which is also proportional to the mean-square error of a non-epistatic model fit to these four genotypes.

**Figure 1:**
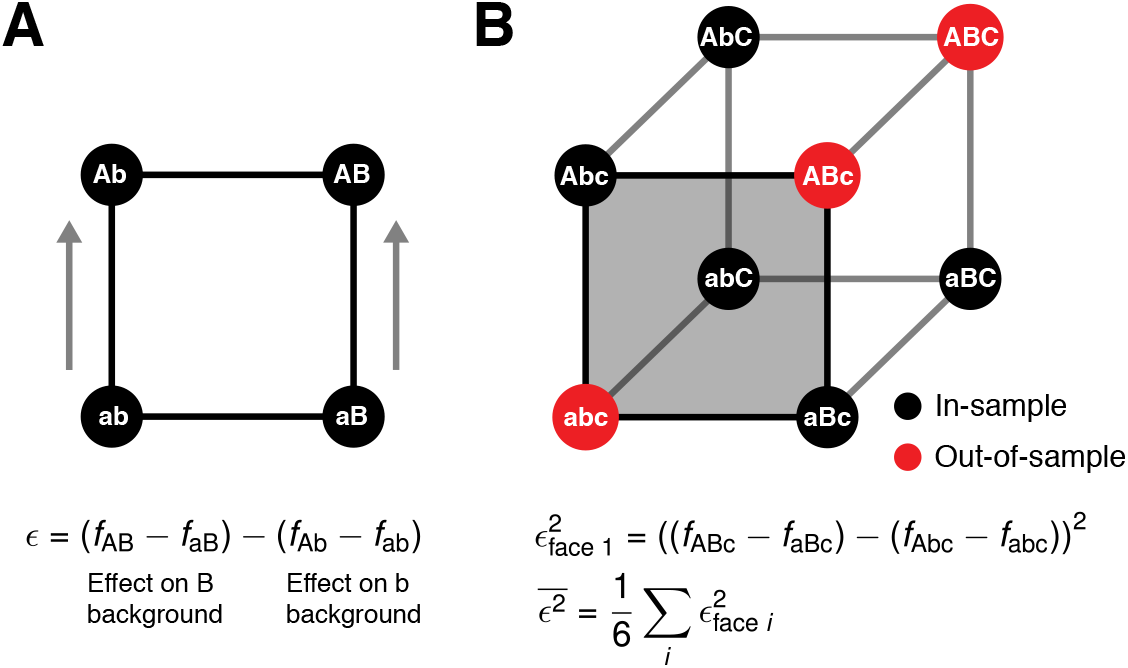
Minimizing average local epistasis. (A) The classical epistatic coefficient *ϵ* measures the difference in the effects of a mutation between two adjacent genetic backgrounds. Here *ϵ* is shown as the difference between the effect of an a → A mutation on a *B* versus *b* background. (B) Larger spaces of genotypes can be decomposed into faces consisting of a WT sequence, two single mutants and a double mutant; one such face is highlighted in gray. For each face, we quantify epistasis locally by calculating the corresponding value of *ϵ*^2^. We then quantify the total amount of epistasis for the sequence-function relationship by taking the average of these values across all faces, 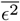. By assigning phenotypic values for the out-of-sample genotypes to minimize 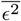, we infer the least epistatic sequence-function relationship compatible with the data in the sense that the average squared difference in the effects of mutations across adjacent genetic backgrounds is as small as possible.

Since we are trying to predict the phenotype for genotype AB by minimizing the change in the effects of each mutation across genetic backgrounds, we can do so by minimizing *ϵ*^2^, giving the solution 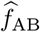:

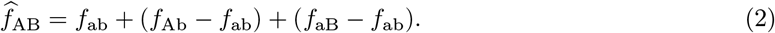

Indeed for this simple case, the effects of mutations can be chosen to be constant across genetic backgrounds, yielding *ϵ* = 0, so that 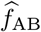 is also the solution given by a traditional non-epistatic model where the effects of mutations are additive across loci.

How can we generalize this solution to larger sequence spaces and more complex patterns of missing data? The traditional choice would be to maintain the additivity of the solution across loci and thus to use a non-epistatic model in the more general case. This is the method biologists have used since Fisher [49]. Here, we take an alternative approach based on minimizing *ϵ*^2^, or rather the expected value of *ϵ*^2^ over all possible pairs of mutations and genetic backgrounds, without imposing additivity or other constraints on the form of epistasis.

More precisely, for genotypes with *l* sites and *α* possible alleles at each site, we can consider the space of possible sequences as a (generalized) hypercube or Hamming graph with 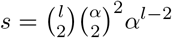 “faces”, each of which consists of 4 genotypes that can be described as a WT sequence together with two single mutants and a double mutant. Any vector **f** that assigns phenotypes to the *α^l^* possible genotypes also defines a value of *ϵ*^2^ for each of these faces, and we denote the average value of *ϵ*^2^ over all such faces as 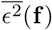, Figure 1B.

Thus, to generalize our solution for the two-locus bi-allelic case with one missing genotype to larger sequence spaces and arbitrary geometric arrangements of the missing data, we want to find the value of **f** that matches our observed phenotypes where available but otherwise minimizes 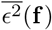. To do this, we note that 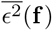 is non-negative, since the *ϵ*^2^ for each face is non-negative, and that the formula for *ϵ*^2^ for each face is a second-degree polynomial and thus so is 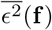. As a result, our constrained minimization problem is in fact a positive semi-definite quadratic minimization problem with an equality constraint, a form of problem that has an analytical solution [50] based on solving a single set of coupled linear equations (see *Materials and Methods*).

In particular, if we write the set of known genotypes as *B* and the minimum epistasis interpolation solution as 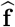, then we first assign 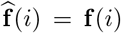 for *i* ∈ *B* to satisfy the constraint that our solution is equal to the observed phenotypic value when available. Then, for each *i* ∉ *B*, the minimum epistasis reconstruction of the sequence-function relationship is given by setting 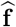 to the solution of the following *α^l^* − |*B*| equations (one equation for each *i* ∉ *B*, see *Materials and Methods*):

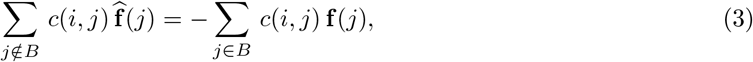

where the values of *c*(*i, j*) depend only on the Hamming distance between genotypes *i* and *j* and are given by:

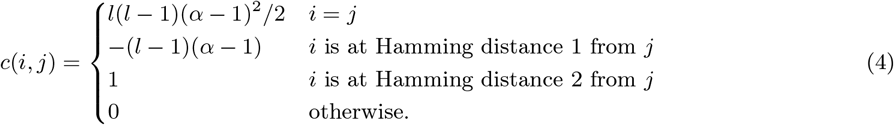

Moreover, these equations have a unique solution if and only if the least squares fit of the corresponding non-epistatic model has a unique solution (see *SI Appendix*).

## Properties of the interpolation solution

Because of its mathematical simplicity, we can in fact provide several guarantees for the properties of this minimum epistasis interpolation solution.

First, consider some focal genotype *i*. This genotype is a member of 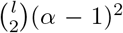 faces, and the phenotypes of the three other genotypes in each face can be used to derive a non-epistatic prediction for the phenotype at *i*. Since these predictions are not necessarily all the same, we can take their mean to produce the average local non-epistatic prediction for genotype *i*. Perhaps surprisingly, the solution to our constrained minimization problem 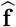 has the property that for any missing genotype *i*, 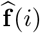 is exactly equal to this local non-epistatic prediction.

Second, the above result can be reinterpreted in geometric terms based on the mean phenotype among genotypes at distance *d* from the focal genotype *i*. Letting **d**_*k*_(*i*) denote the mean value of 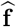 for sequences at distance *k* to *i*, we have

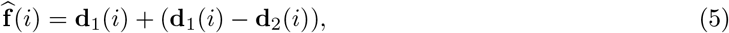

This is similar to a Taylor approximation around i where we correct the nearest-neighbor estimate 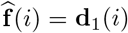 by the difference **d**_1_(*i*) − **d**_2_(*i*), which captures the average effects of the mutations carried by *i* when introduced on mutationally adjacent genetic backgrounds.

Third, the solution has an illuminating connection with the discretized heat equation and the (second-order) Laplace operator. In particular, if **L** is the discrete analog to the continuous Laplace operator (i.e. **L** is the graph Laplacian for our Hamming graph), then our constrained minimization problem is equivalent to a boundary value problem for the second-order discrete Laplace operator **L**^2^ − *α***L**. Interestingly, the solutions to boundary value problems in continuous space for the squared Laplace operator (i.e. the biharmonic equation) are given by the thin plate splines [51], which are widely used in geometric morphometrics [52]. Thus, in many ways it is helpful to think about minimum epistasis interpolation as being a discrete analog of thin plate splines adapted for use in sequence space.

Finally, while our interpolation procedure on its own leaves the observed phenotypes unaltered, it is often useful to also apply some sort of smoothing to the observed data, with the idea of filtering out experimental noise and simplifying the sequence-function relationship to reveal its major features. Our above observations in fact suggest a natural smoothing operator, **M** (see *Materials and Methods*), where applying **M** to a function **f** replaces the value of every sequence with its average local non-epistatic prediction. The key feature of this particular smoothing operator is that applying **M** to 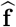 leaves our out-of-sample predictions unaltered. Thus, we can choose to apply **M** to 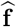 if we prefer to smooth the in-sample data, or work directly with 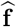 if we prefer to leave the in-sample data unaltered.

## Application to the crater landscape model of transcriptional regulation

To provide a simple demonstration of our interpolation technique, we analyze a biophysically inspired model for transcriptional regulation known as the crater landscape [46]. The model treats a single transcription factor binding site, where the fitness of the binding site is a function of the number of mismatches from the best binding sequence, and where the fitness maximum is achieved at an intermediate distance from the best binding sequence due to selection against spurious binding when the transcription factor is at a low concentration [46] (see *Materials and Methods*). However, for our purposes the most important feature of the model is that it produces a complex pattern of epistasis that can nonetheless be displayed graphically in one dimension (Figure 2), which is helpful in getting an intuitive feeling for the behavior and characteristics of our interpolation procedure.

**Figure 2:**
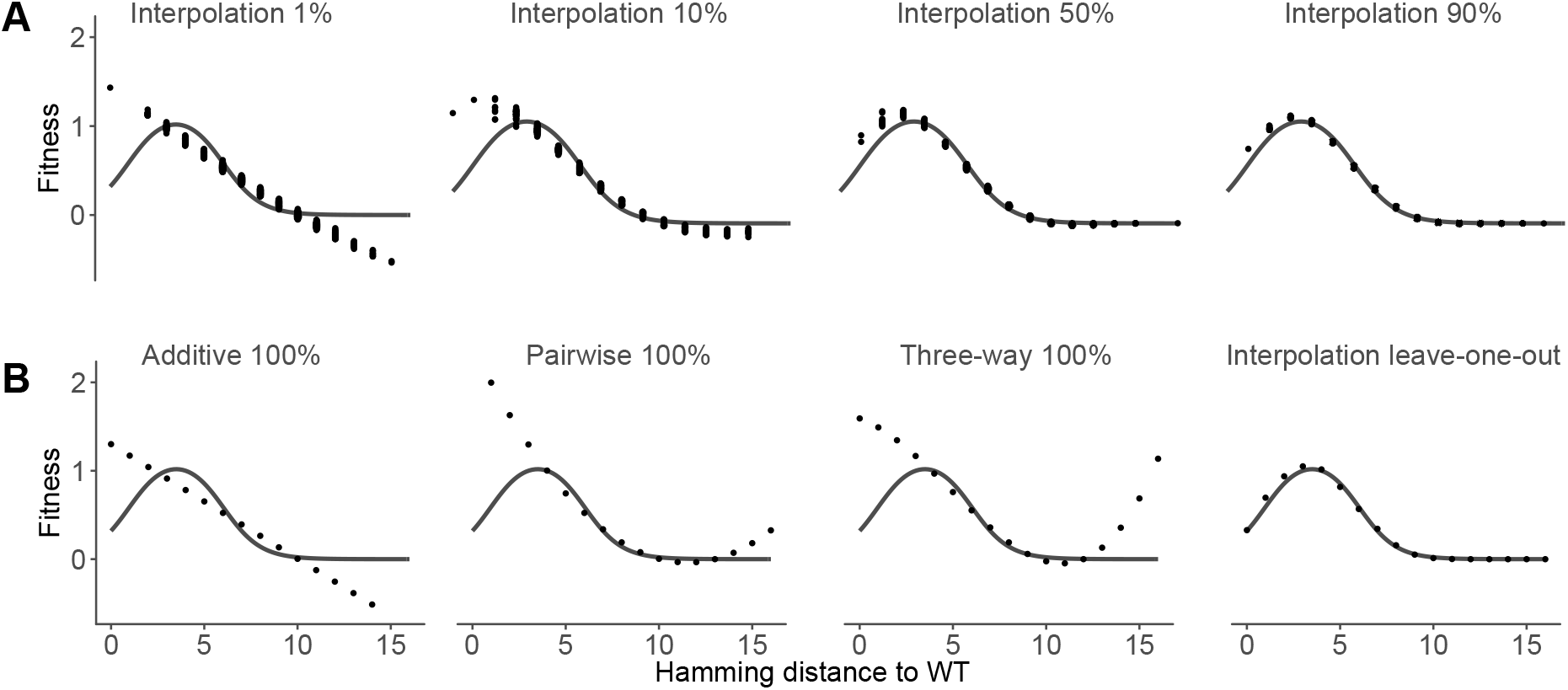
Minimum epistasis interpolation but not low-order regression models can learn the crater model for transcriptional regulation. The crater model produces a fitness landscape where fitness depends only on the Hamming distance to the WT sequence, with an optimum at an intermediate Hamming distance (*l* = 16 and *α* = 2; see *Materials and Methods* for other parameters). Gray curve shows the true fitness landscape. (A) Out-of-sample predictions of the minimum epistasis interpolation with random subsets of 1%, 10%, 50%, and 90% of genotypes used for training. The predictions adapt to the shape of the crater landscape with increasing data density. (B) Reconstruction of the crater landscape by the additive, pairwise, and three-way regression models fitted using ordinary least square with 100% of the data. The interpolation panel shows leave-one-out results (equivalent to applying the smoother **M** to the full landscape).

We first consider the out-of-sample behavior of our interpolation procedure as a function of the fraction of genotypes whose phenotypes are known (Figure 2A). We see that the complexity of the model changes adaptively with the sampling density, producing essentially additive predictions when given the phenotypes for a random 1% of genotypes, but providing an increasingly close fit as the amount of training data increases.

Next, we compare our method to three commonly used regression models, namely the additive model, which assumes independent contribution of sites to fitness, and pairwise and three-way interaction models. The regression models were fit using ordinary least squares with 100% of the data in order to examine the best reconstruction of the true landscape given their respective model complexity. Because our interpolation procedure leaves the data points unchanged, as a fair comparison we make leave-one-out predictions for genotypes of each distance class by giving our method all but one genotype as training data, which is equivalent to smoothing the complete landscape using our smoother M. We find that while the interpolation model can provide a very good fit to this landscape, these lower-order regression models are incapable of producing a qualitatively correct approximation of the landscape, even when given 100% of the landscape as data. This occurs because the crater landscape contains interactions of all orders, and thus cannot be captured by these lower-order interaction models.

## Application to protein G

To test our method on experimental data, we now turn to a combinatorial mutagenesis study of the IgG-binding domain of streptococcal protein G (GB1) [34], a model system for studying protein folding stability and binding affinity [5, 34, 53, 54]. By sequencing a library of protein variants before and after binding to IgG-Fc beads, this experiment [34] attempted to assay all possible combinations of mutations at four sites (V39, D40, G41, and V54; 20^4^ = 160000 protein variants) that had previously been shown to harbor a particularly strong and complex pattern of genetic interactions [5]. Binding scores were determined as log enrichment ratios (logarithm of ratio of counts before and after selection, normalized by subtracting the log ratio of the wild-type), however the original authors could not report binding scores for 6.6% of variants due to low coverage in the input library (10 or fewer input reads [34]).

Here, we use this dataset to both predict the phenotypes for these missing sequences and to assess the performance of our method by making predictions for randomly sampled held-out data. In addition to minimum epistasis interpolation, for comparison we also fit an additive model using ordinary least squares and *L*^2^-regularized pairwise and three-way regression models [26] with regularization parameters chosen by 10-fold cross-validation (see *Materials and Methods*).

We first compare the predictive power of the four models by plotting the out-of-sample *R*^2^ against training sample size, Figure 3A. The three epistatic models substantially outperform the additive model, consistent with the high degree of epistasis previously observed for this dataset. While the pairwise model produces a good fit with relatively little training data and is the best performing model when training data on less than 40% of genotypes is available, its out-of-sample *R*^2^ saturates at 0.78 and fails to improve beyond 20% training data. In contrast, the out-of-sample *R*^2^ for the three-way model and our interpolation method continue to improve and surpass the pairwise model at high data density, indicating the presence of higher-order epistasis in this dataset. Overall, the predictive power of our method and the three-way model were very similar throughout the whole range of sampling, with the interpolation model having marginally better predictive power at low data density and the three-way model performing marginally better at high training data density (test-set *R*^2^ of 0.831 for interpolation and 0.843 for regularized three-way regression at the largest fraction of training data, 92.4%, with the remaining 93.4%-92.4%=1% of observed genotypes reserved as a test set).

**Figure 3:**
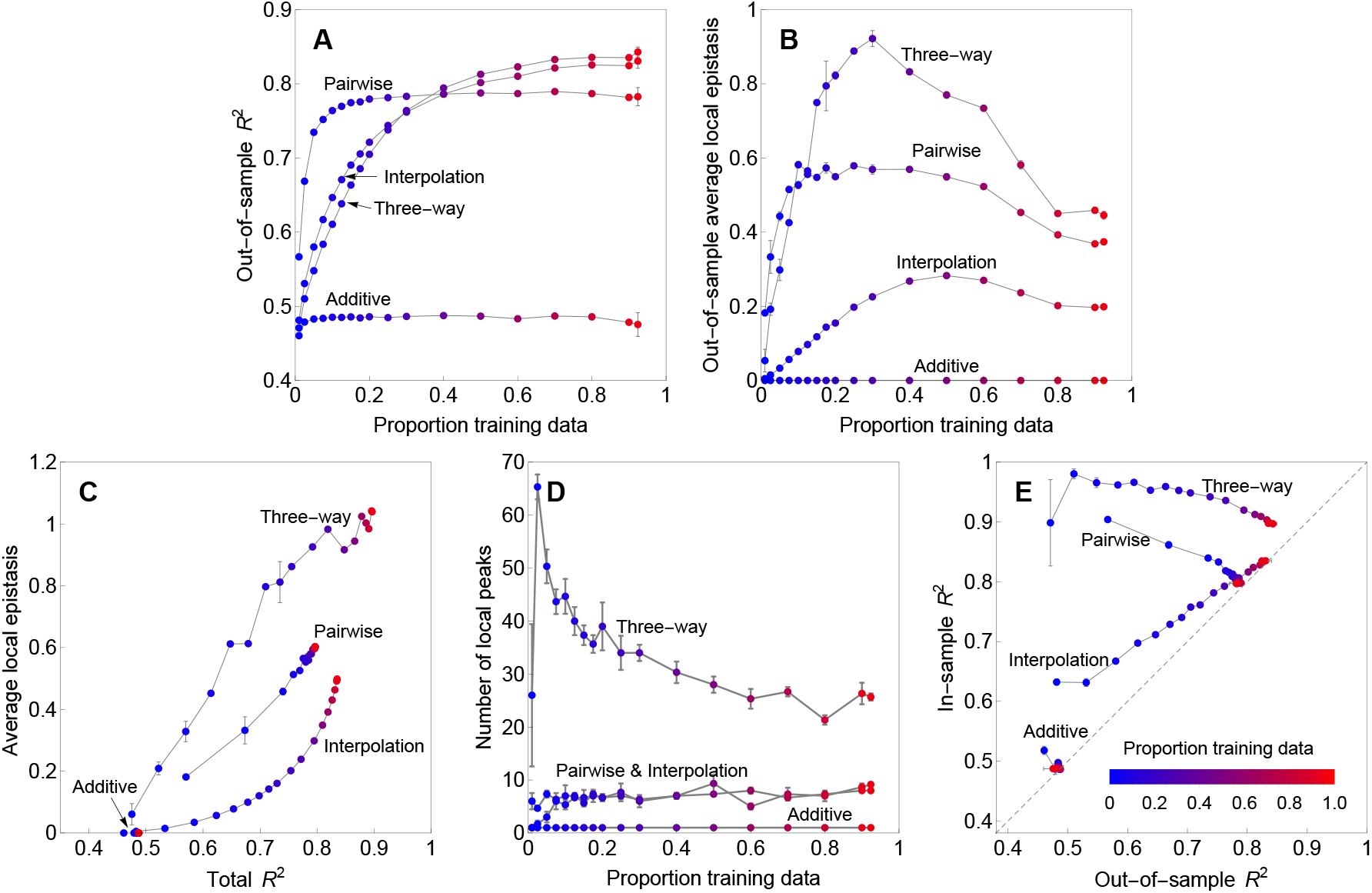
Model performance for the GB1 combinatorial mutagenesis dataset. Additive models were fit using ordinary least squares. Pairwise and three-way interaction models were fit using regularized regression with regularization parameters chosen by 10-fold cross-validation (see *Materials and Methods*). Points are color-coded to represent the proportion of data randomly assigned as training. Error bars indicate one standard error. A) Predictive power (out-of-sample *R*^2^) as a function of the proportion of in-sample genotypes. B) Mean squared epistasis coefficients between random pairs of mutations connecting out-of-sample genotypes as a function of the proportion of in-sample genotypes. C) Global features (in-sample and out-of-sample) of the four models assessed using *R*^2^ between the fitted model and the complete dataset (total *R*^2^) and average local epistasis 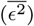. Each model is represented by a curve with points corresponding to increasing proportion of the total dataset assigned as training data. Note that the additive model appears at the lower left part of the plot as its total *R*^2^ quickly stabilizes and its 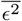 is zero by definition. D) Number of local maxima of the reconstructed landscapes at different training data sizes. The curves of the pairwise model and interpolation exhibit similar trends and therefore are labeled together. E) Model optimism assessed by plotting in-sample *R*^2^ vs. out-of-sample *R*^2^.

However, despite the similar predictive power of the three-way and interpolation models, the interpolation achieves this predictive power using far less epistasis than the three-way model. In particular, Figure 3B shows the mean squared epistatic coefficients between pairs of mutations within the missing and held out data (i.e. mean squared epistasis across all faces contained in the missing and held-out data). We see that the mean squared epistatic coefficient for minimum epistasis interpolation is less than half of the mean squared epistatic coefficient for the three-way model across the whole range of sampling densities and that the interpolation model even has less epistasis than the pairwise model. Overall, we conclude that the predictive power of the interpolation model is quite similar to the three-way interaction model, but that the reconstruction given by the interpolation ought to be preferred because it is far smoother and hence more parsimonious.

So far we have concentrated on out-of-sample prediction, but it is sometimes also useful to consider model behavior within the data in order to reduce the effects of experimental noise and to better reveal the large-scale features of the sequence function relationship. While the three regression models naturally also provide smoothed predictions within the sample, for our interpolation model we first predict all missing data and then apply the smoother **M** which leaves the out-of-sample predictions unchanged while replacing each in-sample observation by the average of the local non-epistatic predictions (i.e. for each genotype, we consider the non-epistatic prediction based on each possible pair of single mutations and the corresponding double mutant, and then replace its observed or inferred value with the average of these predictions).

To examine the global characteristics of these smoothed landscapes we first represented each model as a curve with points corresponding to different training data sizes, Figure 3C, plotting both the *R*^2^ between the fitted model and the complete dataset (total *R*^2^) as a measure of goodness of fit and use the average squared epistasis 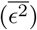 as a measure of the ruggedness of the fitted landscapes. We see that the global behavior of the interpolation model is again quite different from that of the three-way and pairwise interaction models, and at high sampling the smoothed fit deviates from the observed data to an extent that is intermediate between these two regression models, but which is far less rugged than either of them. To provide a different view on the type of epistasis that is incorporated into these smoothed landscapes we also considered the number of local maxima in the reconstructions (Figure 3D). Here, our method constructed landscapes with similar number of local maxima as the pairwise model, while the three-way model produced landscapes with at least three times as many local maxima, again suggesting that our smoothed landscape is providing a simpler reconstruction than the three-way interaction model.

Finally, for the purposes of understanding the qualitative features of the sequence-function relationship, it is desirable that the characteristics of the smoothed landscape are similar across the observed and imputed data, so that any patterns detected correspond to true qualitative features of the sequence function-relationship rather than artifacts due to the pattern of missing data. To evaluate the extent of the consistency between the in-sample and out-of-sample regions of sequence space, we first considered the phenomenon of model optimism [55], where the in-sample *R*^2^ of a fit model can be far higher than its out-of-sample *R*^2^ (Figure 3E). We see that while the three-way and pairwise models have in-sample *R*^2^ that are roughly constant in sampling density and often far higher than the out-of-sample *R*^2^, our smoothed landscape provides a global fit wherein the in-sample and out-of-sample *R*^2^ are well-calibrated to each other, so that the goodness of fit is roughly constant across all of sequence-space.

While Figure 3E shows that the extent of model optimism for the pairwise and three-way interaction models is largely alleviated at high data densities, we paradoxically observed anti-conservative behavior for these models in the high data regime. In particular, when a large fraction of possible genotypes are used as training data, these models appear to suffer from an artifact wherein they have a tendency to predict local maxima at out-of-sample sequences, with this enrichment reaching greater than 3-fold when using our largest fraction of sequence-space for training (92.4% of genotypes, *SI Appendix* Figure 1). In contrast, minimum epistasis interpolation does not exhibit this enrichment, and rather behaves conservatively, showing a depletion of out-of-sample predicted local maxima in the data dense regime (*SI Appendix* Figure 1). Because in studies of sequence-function relationships we are often particularly interested in the positions of these local maxima (e.g. “fitness peaks”), the conservative behavior of minimum epistasis interpolation may be desirable in order to limit the number and frequency of false-positive predictions.

### Structure of epistasis in protein G

We have shown that minimum epistasis interpolation combined with the smoother **M** has the tendency to remove experimental noise and spurious maxima while preserving the large-scale structure of the landscape and accommodating complex higher-order epistasis. This suggests that such methods may also be useful for the interpretation, exploration, and intuitive explanation of empirical data for specific sequence-function relationships. In this section, we combine imputation using minimum epistasis interpolation and the corresponding smoother **M** with a visualization technique developed in [56] to perform exploratory data analysis on the full 20^4^ = 160, 000 genotype GB1 binding landscape [34]. Problems with missing data and a proliferation of noise-driven local optima had previously impeded successful application of this visualization technique to empirical data. We show that the methods used here alleviate these difficulties, allowing for a simple and intuitive analysis of this highly epistatic sequence-function mapping.

In particular, the visualization technique is based on using the GB1 data to construct a model of molecular evolution for these four amino acid positions and creates a low-dimensional representation of the corresponding sequence space that optimally approximates the time for a population to evolve from from one genotype to another under selection for high binding (see *Materials and Methods*). The result is a plot where high-binding (i.e. high-fitness) sequences are broadly separated when it would typically take a long time to evolve from one to the other. Figure 4 shows this visualization for GB1, and indicates that there are three relatively distinct sets of high-binding sequences (warm colors) that would take a long time to evolve from one to the other. These regions contain the vast majority of high-binding sequences (97.5% of sequences with smoothed fitnesses greater than WT and all of the top 100 measured binders are contained within the boxed regions) and appear as protrusions from a core of low-binding sequences (cool colors), plotted near the origin. The figure marks local maxima with black rings, and we see that each of these separate regions of high-binding sequences corresponds to a cluster of one or more local fitness maxima, with the wild-type sequence observed near one of these clusters (WT marked with gray ring).

**Figure 4:**
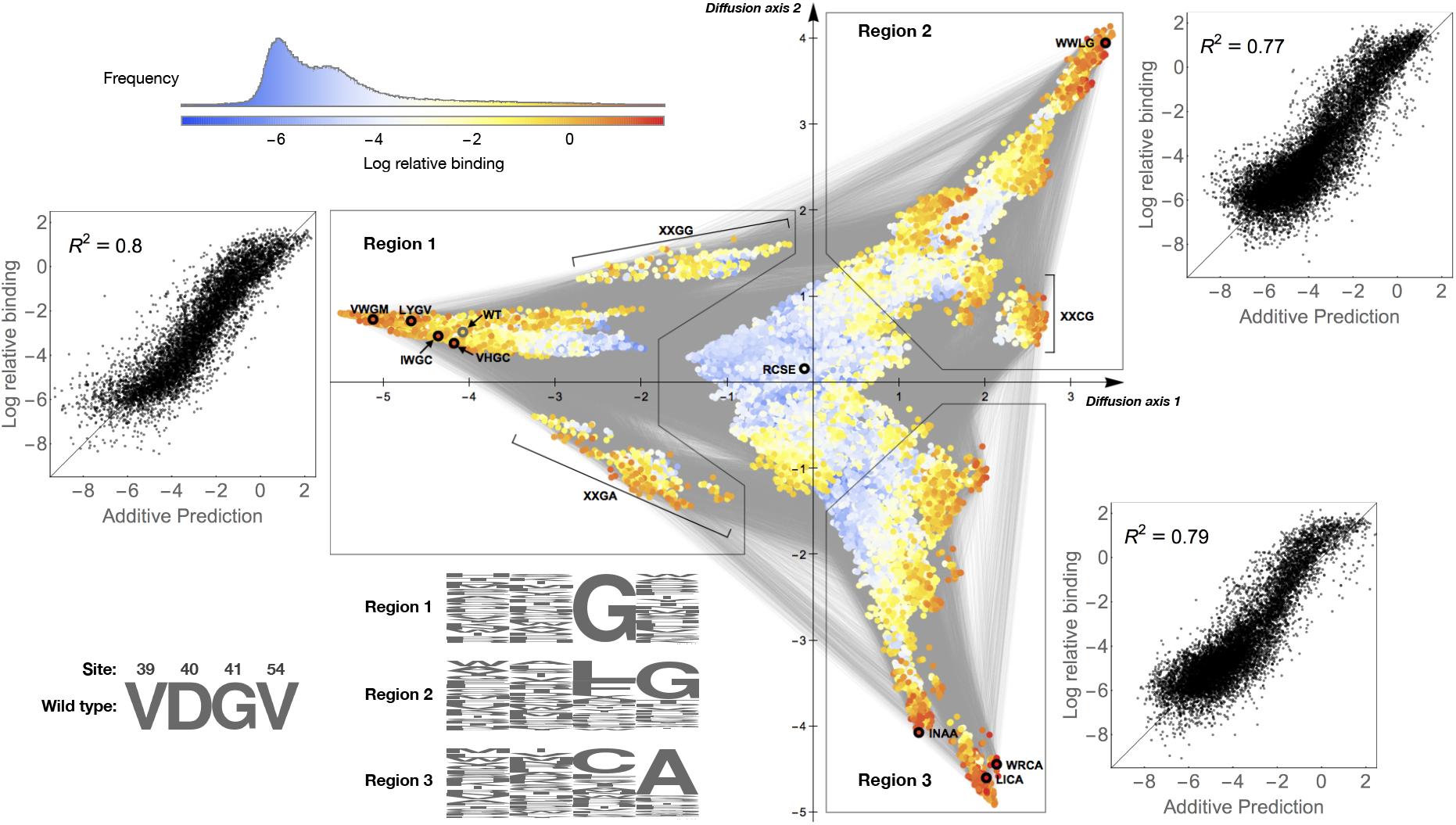
Visualization of the GB1 landscape reconstructed using minimum epistasis interpolation and the local non-epistatic smoother. Genotypes are plotted using the dimensionality reduction technique from [56] (see *Materials and Methods*). Points are genotypes, colored according to their smoothed binding phenotype, and two genotypes are connected by an edge if they differ by a single amino acid substitution. Local fitness peaks are highlighted by black circles. The x and y axis are, respectively, the first and second diffusion coordinate and have units of square-root expected neutral substitutions per site. Three high fitness regions are characterized by their distinct sequence composition (sequence logos). The scatter plots show the fit of an additive model to the unsmoothed binding values within each of the three high-binding regions. These scatter plots indicate that, despite the complex pattern of epistasis in the landscape as a whole, the sequence-function relationship is approximately additive within each individual high-binding region.

To better understand the characteristics of these three high-binding regions and their underlying biophysical explanation, we constructed sequence logos to identify the common features of the sequences within each region (Figure 4). We see that the key characteristic of the first region, which contains the wild-type sequence, is that they all have a glycine at position 41. However, the composition of the other two regions is more complex. Sequences in region 2 often have a glycine at position 54 instead of 41, and the highest binders tend to instead have a leucine or phenylalanine at site 41. In region 3, sequences typically have an alanine instead of a glycine at site 54, with the highest binders generally having either a cysteine of another alanine at site 41. At a more biophysical level, sites 41 and 54 directly interact in the GB1 crystal structure, albeit as part of dynamically active and conformationally variable portion of the protein [5, 34]. In particular, [5, 34] previously suggested that the epistasis observed at these sites was in part due to steric interactions between sites 41 and 54. This analysis is consistent with our observation that the major differences between the three high-binding regions lie in the identity (glycine or alanine) and placement (site 41 or 54) of the small non-polar residue relative to bulkier amino acids (e.g. leucine, phenylalanine or cysteine at site 41).

Finally, we considered the structure of the fitness landscape within each of the high-binding regions. Perhaps surprisingly, we found that within each region even the unsmoothed values are reasonably approximated by a simple additive model (scatter plots in Figure 4, *R*^2^ of .77 to .8, randomization test *p* < 0.003 for each region, see *Materials and Methods*). However, these models differ substantially between the three regions, particularly with respect to the additive effects of substitutions at sites 41 and 54 (*SI Appendix* Fig 2), and all these models fit substantially better than the global additive model investigated previously (*R*^2^ = 0.49), indicating that overall the sequence-function relationship appears to be locally rather than globally additive.

In summary, we are left with a qualitative understanding of the structure of this fitness landscape at several different levels of detail. At the coarsest level, we find that although the GB1 fitness landscape harbors a substantial degree of epistasis, in large part this arises from the presence of three distinct high-fitness regions, and that the fitness landscape is approximately additive within each such region. At a finer level of detail, we observe the presence of multiple local fitness maxima within some of these regions. Finally, our visualization (Figure 4) provides a rich depiction of the finer-scale structure of the landscape, suggesting many hypotheses that are ripe for further exploration. For instance, the visualizations show what appear to be fitness “ridges” connecting one high-binding region to another (e.g. the XXGG sequences connecting Region 1 to Region 2 and the XXGA sequences connecting Region 1 to Region 3, respectively), that can serve as paths of moderate fitness that a population might be most likely to take when traversing from one high-fitness region to another. Importantly, these insights all depend on the application of our smoother, which simplifies the landscape by removing features attributable to experimental noise and fully random epistasis in order to reveal its large-scale features.

## Discussion

Understanding the mapping from genotype to phenotype is a key problem for much of biology, from applied areas such as protein design [44, 57], antigenic evolution [58], and the emergence of drug resistance [59], to more basic questions about the repeatability of adaptation [60] and the dynamics of long-term molecular evolution [31]. While the astronomical number of possible genotypes may put a fully comprehensive understanding of this mapping forever out of reach, modern high-throughput experiments are currently providing phenotypic measurements for tens of thousands to millions of genotypes at a time, so that there is a need for computational techniques to translate these high-throughput measurements into phenotypic predictions for genotypes that have not yet been assayed. Here we have presented a principled and highly conservative solution to this problem by inferring the least epistatic possible sequence-function relationship compatible with the observed data in the sense that mutational effects change as little as possible between mutationally adjacent genetic backgrounds while exactly matching the data where available.

One simple way of understanding our approach is by contrasting it with the classical non-epistatic model [49], since both models in some sense minimize the amount of epistasis but do so in different ways. In a non-epistatic model, one assumes that the sequence-function relationship is completely additive so that the effects of mutations are constant across genetic backgrounds and, consequently, the mean-squared epistatic coefficient between random pairs of mutations across random backgrounds is constrained to be precisely zero. One then determines these mutational effects by minimizing the mean squared error of the model predictions for genotypes where data is available.

In minimum epistasis interpolation, these choices are exactly reversed. Whereas a non-epistatic model minimizes the mean square error under the constraint that the mean square epistatic coefficient is precisely zero, here we constrain the reconstruction to exactly match the data so that the mean square error is precisely zero and infer the missing values by minimizing the mean square epistatic coefficient. This allows the data itself to dictate the amount and character of epistasis that is included, since the reconstruction is as additive as possible while still being highly epistatic in regions of sequence space where the data requires it. (In the *SI Appendix*, we show that the classical non-epistatic model and minimum epistasis interpolation can actually be viewed as two ends of a continuum of models that minimize a convex combination of mean-square error and mean-square epistasis, and which all have out-of-sample properties similar to minimum epistasis interpolation).

Our method also provides new insights into the interpretation of higher-order epistatic interactions, that is interactions between mutations at three or more sites. When viewing genotype-phenotype mappings from a regression or analysis of variance standpoint, there is a tendency—going back to the very earliest days of statistics and experimental design [61, 62] —to assume that higher-order interaction terms are likely to be small (e.g. in partial factorial designs where they are purposefully confounded with main effects and lower-order interactions, [63]). However, there is a growing consensus that such higher-order interactions are not only common in genotype-phenotype maps [10, 18, 29, 32, 38] but are expected even for very simple, smooth genotype-phenotype relationships such as where the observed phenotype is just an additive trait that has been run through a nonlinear transformation [31, 32, 40, 64–66]. Our results contribute to this view by showing that the incorporation of higher-order interactions in fact allows substantially less epistatic fits than standard pairwise models. To see why this is the case, it is helpful to realize that higher-order genetic interactions can be thought of as pairwise interactions whose strength changes over different regions of sequence space, which in particular allows the strength of pairwise epistatic interactions to decay towards zero in regions of sequence space that are data-poor or where the interaction is not supported.

Besides viewing genotype-phenotype maps as being defined by sums of interactions between sites as in regression models [26, 29, 67], there are a rich variety of other formalisms for describing genetic variation that are related to the techniques we have developed here [68–72]. Probably the most relevant of these is the correlation between the effects of mutations measured in mutationally adjacent genetic backgrounds, *γ* [10, 71]. Conceptually, maximizing *γ* would be quite similar to our method except that *γ* depends on both *ϵ*^2^ and the variance in the phenotypic effects of mutations [71], so that maximizing *γ* would tend to inflate the magnitude of mutational effects, in essence minimizing the relative rather than absolute amount of epistasis. Our face-specific epistatic coefficients are also related to the “circuit” approach of [69] in that these epistatic coefficients correspond to a subset of the possible circuits (specifically those corresponding to conditional epistasis). However, at a deeper level our approach is most closely related to the Walsh-Fourier decomposition [29, 67, 73–75], where the phenotype is expanded in terms of the eigenvectors of the graph Laplacian **L**, which are also the eigenvectors of the second-order discrete Laplace operator **L**^2^ − *α***L**, so that our minimization problem can be re-cast as minimizing a weighted sum of squared Walsh coefficients, where the weight increases quadratically with interaction order (see *SI Appendix*). Finally, while boundary-value problems involving the graph Laplacian **L** arise in many areas of applied mathematics [e.g. 76–78], here we are faced with a more unusual boundary value problem for **L^2^** − *α***L**. Interestingly, this second-order character arises because of our stipulation that the mutational effects—rather than the phenotypic values themselves—change smoothly as we move through sequence space, whereas naive interpolation based on **L** results in an unrealistic degree of sign epistasis where e.g. multiple deleterious mutations combine to be the average of the single mutations rather than their sum.

While our interpolation procedure exactly matches the data where available, some degree of smoothing is often helpful to better understand the large-scale features of the sequence-function relationship and to ameliorate the effects of experimental noise. To address this need, we proposed a smoother that is philosophically similar to LOESS [79] in that it approximates the sequence-function relationship as being locally additive while making no assumptions about its global structure. Specifically, the smoother replaces the phenotypic value for each genotype with the average of the non-epistatic predictions that would be obtained by taking each possible double mutant as the wild-type. Because of the large number of single and double mutants that this smoothed estimate averages over, such smoothing greatly decreases the impact of experimental noise (e.g. using Equation 5, we can derive the crude estimate of a *l*(*α* − 1)/5-fold variance reduction under IID noise, which for the GB1 study comes out to a roughly 15-fold reduction). Moreover, because the smoother does not alter the out-of-sample predictions given by minimum epistasis interpolation, we see that the interpolation predictions are similarly insensitive to experimental noise. It is important to note that by the same argument the application of the smoother will largely remove any true fully random component of the sequence-function relationship (i.e. the so-called house-of cards component [30, 40, 80]). Thus, for applications where we are most interested in the genotype with high phenotype values (high fitness or highly functional genotypes), concordance between the smoothed and raw experimental phenotypes for a high functionality provides confidence not only that the genotype is likely to be truly functional, but that this functionality is due to a consistent tendency in the local sequence-function relationship rather than some idiosyncratic feature of the individual genotype.

Minimum epistasis interpolation provides a principled and highly conservative method for reconstructing sequence-function relationships that has no tunable parameters and allows epistatic interactions of all orders. Nonetheless, the method has two main sets of limitations that will require additional work to address. First, as implemented here, the computational complexity of the method scales with the number of unobserved genotypes. While in the *SI Appendix* we describe a kernelized implementation whose complexity scales with the number of observed rather than unobserved genotypes, practically these two implementations can only analyze studies whose genotypic spaces are not much larger than the size of the GB1 dataset (160,000 genotypes) or, alternatively, studies containing less than roughly ten thousand observed sequences. Second, we emphasize that despite its many interesting and useful properties, the method introduced here produces only the least epistatic possible reconstruction of the sequence-function relationship and hence is almost necessarily underfitting the data. More general statistical approaches that better reflect the character of epistasis found in a specific dataset are likely possible and capable of providing better out-of-sample performance.

## Materials and Methods

### Formulation of the minimization problem and its solution

Suppose our sequence space consists of two bi-allelic loci (*α* = 2, *l* = 2) and hence four possible genotypes {ab, aB, Ab, AB}. Given a vector that assigns phenotypes to all four genotypes, **f**^*T*^ = [*f*_ab_ *f*_aB_ *f*_Ab_ *f*_AB_]^*T*^, we can calculate the squared epistatic coefficient for **f** as

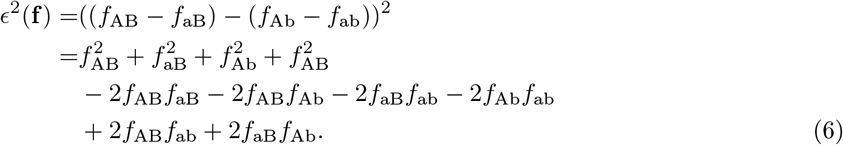

Extending this to arbitrary *α* and *l*, for a vector **f** defined on the set of all *α^l^* genotypes we can calculate the mean squared epistatic coefficient 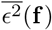 by averaging over all 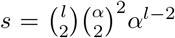 faces of the sequence space. This results in a positive semi-definite quadratic form 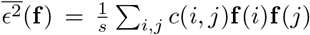, where the *c*(*i, j*) can be found by counting the number of times the ordered product **f**(*i*)**f**(*j*) appears when summing Equation 6 over all faces and which only depend on the Hamming distance *d*(*i, j*) between sequences *i* and *j*. First the squared term for any given genotype (distance 0) appears in 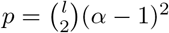 faces with coefficient 1. Second, each ordered pair of genotypes that are at Hamming distance 1 from each other appear in (*l* − 1)(*α* − 1) faces with coefficient −1. Third, if the ordered pair of genotypes are separated by distance 2, they appear in exactly one face with coefficient 1. Thus, we arrive at Equation 4 in the main text.

Now, arrange the coefficients *c*(*i, j*) in a matrix **C** with **C**(*i, j*) = *c*(*i, j*)/*s* and 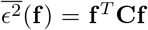. Given data **y** ∈ ℝ^*m*^ for a subset of sequences *B* of size *m* of the set of all possible sequences 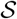, we write 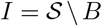 to be the set of all missing sequences. Without loss of generality, we will order our sequences so that the *m* sequences in *B* whose phenotypes are known come first. Our aim is to infer a full landscape 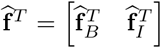 that minimizes the average squared epistatic coefficient under the constraint that we do not change the values for genotypes in *B*. We can formulate this as a quadratic minimization problem with equality constraint:

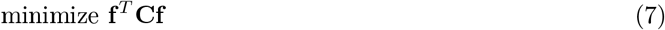

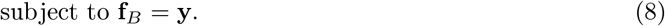

Using *B* and *I* to index submatrices of **C**, we can solve this minimization problem by differentiating **f**^*T*^**Cf** with respect to **f**_*I*_ and setting the gradient to zero. This gives us:

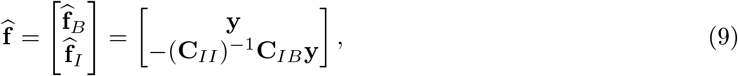

which is equivalent to Equation 3 in the main text (the matrix (**C**_*II*_)^−1^ exists if and only if the non-epistatic model fit by least squares has a unique solution, see *SI Appendix*).

### Mathematical properties of the solution

Rearranging Equation 9 as 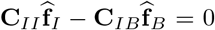, we find the solution 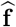 must satisfy 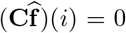, for all genotypes *i* in *I*. To understand what this condition means, we rescale our cost matrix and write it as 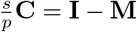, where **I** is the identity matrix. Using the definition of **C** and Equation 4 gives us

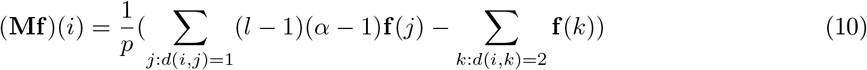

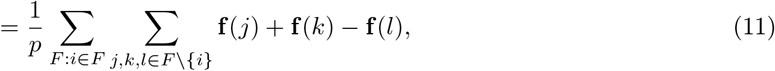

where we enumerate through all *p* faces *F* that *i* belongs to. For each face *F*, **f**(*j*) + **f**(*k*) − **f**(*l*) = **f**(*j*) − **f**(*l*) + **f**(*k*) − **f**(*l*) + **f**(*l*) is the non-epistatic prediction based on sequences *j, k* which are at distance 1 to *i* and *l* which is at distance 2 to *i* (Equation 2). Therefore, (**Mf**)(*i*) returns the average local non-epistatic prediction for *i* based on all faces *i* is in. Thus the necessary condition for the solution 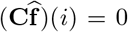 is equivalent to 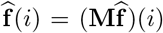, ∀*i* ∈ *I*. That is, for any unknown genotype *i*, its inferred value must be equal to its average local nonepistatic prediction based on 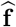. As a result, applying **M** to 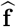 does not alter our predictions for the out-of-sample genotypes.

Let **d**_*k*_(*i*) denote the mean value of 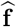 for sequences at distance *k* to *i*. We can rewrite Equation 10 as

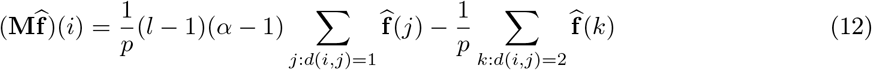

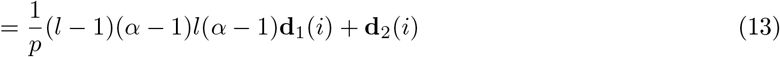

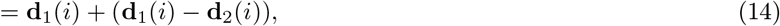

which gives us a geometric interpretation for our method (Equation 5 in the main text).

Our minimization problem also has a close relation to the Dirichlet problem on graphs [78]. To see this, recall the definition of the graph Laplacian **L** for our Hamming graph of all possible sequences

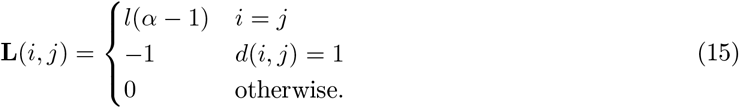

A discrete Dirichlet problem is formulated as finding a function 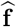 defined on the graph so that 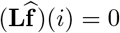 on the unsampled genotypes (interior) *i* ∈ *I*, while satisfying the condition 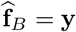.

It turns out that we can re-express **C** in terms of **L** and **L**^2^. In particular for **L**^2^ we have

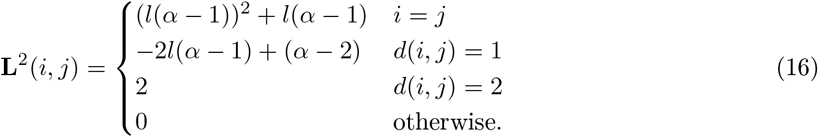

Consequently, **L**^2^ − *α***L** = 2*s***C**. Thus, instead of being harmonic in the interior, i.e. 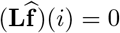, as in the classical discrete Dirichlet problem, the solution to our problem instead must satisfy 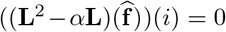 for *i* ∈ *I*, in addition to the boundary value constraint.

### Simulation of crater landscape

We simulated data under the crater landscape model [46] for the fitness of a transcription factor binding site. Specifically, we assume the effects of mutations on binding energy to be constant and the binding probability of any sequence is simply given by its Hamming distance *d* to the best binding sequence: 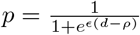, where *ϵ* is the binding energy per nucleotide mismatch and the compound parameter *ϵρ* is the chemical potential measuring the factor concentration [46]. In this minimal model we assume there are two cellular states. The *on* state favors the expression of the gene, hence selects for high binding probability with selection coefficient *s*_on_. The *off* state disfavors gene expression and selects against high binding with coefficient *s*_off_ = −*s*_on_. The total fitness of a sequence at distance *d* is given by

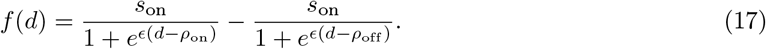

We choose the following parameters *ϵ* = 1, *ρ*_on_ = 6, *ρ*_off_ = 1, and *s*_on_ = 1. We use this model for simulating fitness landscape data for the set of all possible mutants corresponding to a sequence space with *l* = 16 sites and two allelic states at each site.

### *L*_2_ regularized regression

We use the following linear model to fit pairwise and three-way interaction models to the GB1 dataset.

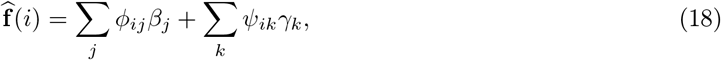

or in matrix notation

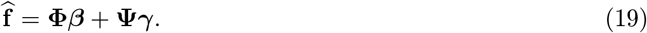

The matrix **Φ** has as columns any orthonormal set of vectors that span the space of non-epistatic fitness landscapes (eignespace of the graph Laplacian **L** associated with eigenvalues 0 and *α*). The columns of **Ψ** form an orthonormal set of vectors that span the space of all pairwise or pairwise and three way functions (eignespace of **L** associated with eigenvalue 2*α* (pairwise) or 2*α* and 3*α* (three-way). see *SI Appendix*).

We fitted *L*_2_ regularized pairwise and three-way regression models to different training data sets *B*. Specifically, we find our solution by minimizing

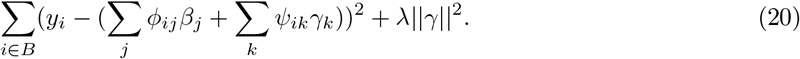

Note that we do not penalize the additive component of the fitted model, to ensure a fair comparison with our interpolation procedure. The regularization parameter *λ* is chosen from a set of potential parameters equally spaced on the log_10_ scale. For each training sample, we performed 10-fold crossvalidation to generate average mean squared errors using the candidate *λ*’s. The *λ* with the least crossvalidated MSE is used to fit the training data and make predictions for the test dataset.

### Visualization of the GB1 landscape

We consider a population evolving in continuous time under weak mutation e.g [81–83] on the full 20^4^ = 160, 000 genotype GB1 binding landscape smoothed using **M**. Specifically, we model evolution as a continuous-time Markov chain where the population moves from genotype to genotype at each fixation event. The rate matrix *Q* of the Markov chain is

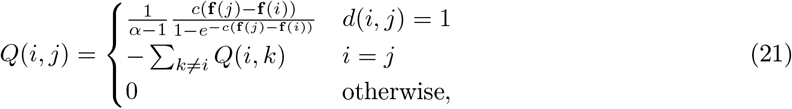

where *c* is the conversion factor that transforms log binding to scaled fitness (Malthusian fitness × *N_e_*). We choose *c* so that the expected log-binding at stationarity is equal to the log-binding of the wild type. Time is scaled so that the mutation rate per site is equal to 1. For a 2-dimensional representation of the GB1 landscape, we use as coordinates the right eigenvectors of *Q* associated with the 2 largest nonzero eigenvalues. This allows our low-dimensional representation of the landscape to optimally capture the expected time for a population to evolve from genotype *i* to *j* [47].

### Significance test of *R*^2^ for local additive fits

To assess the statistical significance of the total *R*^2^’s of additive models fit to the three regions identified in our visualization of the GB1 landscape, we sampled, for each region, 1000 random subsets of the same size. We then fit additive models to these random subsets to calculate the null distribution of total *R*^2^. We calculate the *p*-value for the *R*^2^ for each region as the fraction of random subsets that have equal to or greater *R*^2^ than the observed value.

## Supporting information

Mathematica notebook and data file

## Data Availability

Computer code to replicate all analyses is included in the Supporting Information in the form of a Mathematica notebook.

