## Supplementary material for "Minimum epistasis interpolation for sequence-function relationships": SI appendix

June 1, 2019

#### 1 Mathematical preliminaries and relationship with the Walsh-Fourier approach

In this introductory section of the SI Text we review the Walsh-Fourier approach to epistasis and recast our minimum epistasis interpolation within a Walsh-Fourier framework. We also prove some simple results that will be helpful in the remaining sections of this document.

From a mathematical standpoint, the Walsh-Fourier approach to analyzing epistasis [1–4] is best viewed as a special case of the Fourier analysis of functions on graphs [5], which itself is simply the decomposition of a function on a graph in terms of the eigenvectors of the associated graph Laplacian. However, the key biological facts that gives this particular application its character is that (1) genotypes are combinatorial objects made up of alleles at some set of sites and (2) these genotypes are converted from one to another by mutations that occur at the individual sites. As a result, the graph describing the possible mutational transformations among the set of all possible genotypes takes the form of a product graph. Moreover, because biological sequences are generally composed of $\alpha$ possible states (e.g. 4 nucleotides or 20 amino acids) at each of $l$ sites, the factors in these product graphs are identical, and this results in a graph Laplacian with only $l+1$ distinct eigenvalues and highly degenerate eigenspaces that correspond to the different possible orders of genetic interactions. Importantly, and in contrast to the case for more general (i.e. less symmetrical) graphs, these eigenspaces can be characterized analytically, and this allows formal analysis even when the number of possible genotypes $n = \alpha^l$ is astronomically large.

For a general graph $G$, recall that the associated graph laplacian $\mathbf{L}$ is given by the matrix

$$\mathbf{L}(i,j) = \begin{cases} \text{degree of vertex } i & i = j \\ -1 & i \text{ is adjacent to } j \\ 0 & \text{otherwise.} \end{cases} \tag{1}$$

In addition, since $\mathbf{L}$ is real and symmetric, it admits the eigendecomposition

$$\mathbf{L} = \mathbf{Q}\mathbf{\Lambda}\mathbf{Q}^T = \sum_{i=1}^{n} \lambda_i \mathbf{v}_i (\mathbf{v}_i)^T, \tag{2}$$

where $\mathbf{Q}$ is an orthogonal $n \times n$ matrix whose $i$-th column is the eigenvector $\mathbf{v}_i$ of $\mathbf{L}$, and $\mathbf{\Lambda}$ is the diagonal matrix with the corresponding eigenvalues. If we order the eigenvectors in order of increasing

$\lambda_i$, the corresponding eigenvectors $v_i$ are functions that change increasingly quickly as we move across the graph [6], so that the eigenvalue $\lambda_i$ can be interpreted as the analog of the frequency in the classical Fourier decomposition. Moreover we can decompose any function $f$ on the graph into the contributions corresponding to any given frequency via multiplication by the projection matrices $\mathbf{v}_1(\mathbf{v}_1)^T, \ldots, \mathbf{v}_n(\mathbf{v}_n)^T$.

We now turn to the corresponding eigendecomposition for our specific graph of interest, which is the Hamming graph on words with $l$ letters and an alphabet of size $\alpha$. More biologically, this is the graph of all possible sequences of length $l$ with $\alpha$ possible alleles per site where two sequences are adjacent if and only if they differ a point mutation, that is, their Hamming distance is equal to 1. Define the Cartesian product of two graphs $G$ and $H$, $G \times H$ to be the graph with vertex set $V(G) \times V(H) = \{(u, u') | u \in V(G), u' \in V(H)\}$ where two vertices $(u, u')$ and $(v, v')$ are adjacent on $G \times H$ if and only if either $u = u'$ and $v \sim v'$ or $u \sim u'$ and $v = v'$. Then it is easy to verify that the Hamming graph is the $l$-fold Cartesian product of the complete graph on $\alpha$ vertices [7], with the complete graph corresponding to the genotypic space with only one site.

An important, and easy to verify, result about product graphs is that if $\lambda$ and $\mu$ are Laplacian eigenvalues of graphs $G$ and $H$ with eigenvectors $\mathbf{v}$ and $\mathbf{u}$ respectively then the vector whose entry for $(u, u')$ is $\mathbf{v}(u)\,\mathbf{u}(u')$ is a Laplacian eigenvector of $G \times H$ with eigenvalue $\lambda + \mu$ [8]. As a result, if $\lambda$ and $\mu$ have multiplicities $a$ and $b$ , then $\lambda + \mu$ is a Laplacian eigenvalue of $G \times H$ with multiplicity $ab$. Now, for the complete graph (or equivalently the special case where $l = 1$, it is easy to verify $\mathbf{L}$ has eigenvalues 0 and $\alpha$, with multiplicity 1 and $\alpha - 1$, respectively. Thus it follows that the Laplacian for our genotypic space, which is the $l$-fold product of complete graphs, has only $l + 1$ distinct eigenvalues $\lambda_k = \alpha k$, $k = 0, \ldots, l$ with multiplicity $m_k = \binom{l}{k}(\alpha - 1)^k$ [2]. Let $\{\mathbf{u}_{k_i}\}_{1 \leq i \leq m_k}$ be a set of orthonormal eigenvectors that span the eigenspace associated with $\lambda_k$. We can rewrite Equation 2 as

$$\mathbf{L} = \mathbf{Q}\boldsymbol{\Lambda}\mathbf{Q}^T = \sum_{k=0}^{l} \lambda_k \sum_{i=1}^{m_k} \mathbf{u}_{k_i}(\mathbf{u}_{k_i})^T = \sum_{k=0}^{l} \alpha k \sum_{i=1}^{m_k} \mathbf{u}_{k_i}(\mathbf{u}_{k_i})^T, \tag{3}$$

We can further simplify by setting $\mathbf{W}_k = \sum_{i=1}^{m_k} \mathbf{u}_{k_i}(\mathbf{u}_{k_i})^T$. Thus $\mathbf{W}_k$ is the projection matrix for the $k$-th eigenspace. Importantly, $\mathbf{W}_k(i, j)$ only depends on $k$ and the Hamming distance $d(i, j)$ between the two genotypes $i$ and $j$ and is invariant to the choice of basis and is given by the Krawtchouk polynomial $w_k(d(i, j))$ [3, 9]:

$$w_k(d) = \alpha^{-l} \sum_{q=0}^{\min(k,d)} (-1)^q (\alpha - 1)^{k-q} \binom{d}{q} \binom{l-d}{k-q}. \tag{4}$$

Critically, the $(l + 1)^2$ possible values of this expression for $k = 0, \ldots, l$ and $d = 0, \ldots, l$ can be pre-calculated, which allows efficient entry-wise evaluation of many of the matrices that follow.

As in the general case, given this eigendecomposition we can decompose a function $f$ on the space of genotypes into contributions corresponding to each of these $l + 1$ eigenspaces, where in fact the $k$-th eigenspace corresponds to interactions between $k$ sites in the sequence, i.e. $k$-th order interactions [4], and these components of $f$ can be derived via multiplication by the projection matrices $W_k$. To put this another way, we have a direct sum decomposition $\mathbb{R}^n = V_0 \oplus V_1 \oplus \cdots \oplus V_l$, with $V_k = \{\mathbf{v} : \mathbf{L}\mathbf{v} = \lambda_k \mathbf{v}\}$ being the $k$-th eigenspace. In what follows, we will be particularly interested in the space of non-epistatic functions $V_0 \oplus V_1$. Since $m_k = 1$ and $l(\alpha - 1)$ for $k = 0$ and 1, $\dim(V_0 \oplus V_1) = 1 + l(\alpha - 1)$. If we consider an arbitrary basis for this space, we can put these basis vectors into a matrix $\mathbf{X} \in \mathbb{R}^{n \times (1+(l-1)\alpha)}$, where $n = \alpha^l$, so that any non-epistatic function can be expressed as $\mathbf{X}\boldsymbol{\beta}$ for some choice of $\boldsymbol{\beta}$ (one natural option for this basis is $\mathbf{X} = [\mathbf{1} \quad \mathbf{X}_{\text{additive}}] \in \mathbb{R}^{n \times (1+(l-1)\alpha)}$, where $\mathbf{1}$ is the constant vector of all ones, and $\mathbf{X}_{\text{additive}}$ has $(l - 1)\alpha$ columns, with the $k_i$-th column consisted of an indicator variable encoding the presence or absence for all genotypes of a non-WT state $i$ on site $k$ for some choice of WT genotype, but for our purposes any basis will do). More generally, the epistatic coefficients (Walsh coefficients) are given by the products $b_{k_i} = (\mathbf{u}_{k_i})^T \mathbf{f}$ so that we can represent $\mathbf{f}$ as with respect to the basis $\{\mathbf{u}_{k_i}\}$ as [1, 3]

$$\mathbf{f} = \sum_{k=0}^{l} \sum_{i=1}^{m_k} b_{k_i} \mathbf{u}_{k_i}. \tag{5}$$

With these basic results and notation in hand, we can return to the subject of minimum epistasis interpolation. In the *Materials and Methods* section of the main text we showed that $\mathbf{L}^2 - \alpha \mathbf{L} = 2s\mathbf{C}$, so we can immediately write down the eigendecomposition of $\mathbf{C}$ as

$$\mathbf{C} = \frac{1}{2s} \mathbf{Q} \left( \mathbf{\Lambda}^2 - \alpha \mathbf{\Lambda} \right) \mathbf{Q}^T = \sum_{k=0}^{l} \lambda'_k \sum_{i=1}^{m_k} \mathbf{u}_{k_i} (\mathbf{u}_{k_i})^T, \tag{6}$$

with $\lambda'_k = \frac{\lambda_k^2 - \alpha \lambda_k}{2s} = \frac{\alpha^2}{2s} \left( k^2 - k \right)$. Importantly, this gives us $\lambda'_0 = \lambda'_1 = 0$, which means that the null space of $\mathbf{C}$, $\mathrm{N}(\mathbf{C})$, is identical to the non-epistatic subspace, i.e. $\mathrm{N}(\mathbf{C}) = V_0 \oplus V_1 = \mathrm{col}(\mathbf{X})$.

Using this representation for $\mathbf{C}$, we find

$$\overline{\epsilon^2}(\mathbf{f}) = \mathbf{f}^T \mathbf{C} \mathbf{f} = \sum_{k=0}^{l} \lambda'_k \sum_{i=1}^{m_k} b_{k_i}^2. \tag{7}$$

This shows that our minimization problem can be recast as minimizing a weighted sum of squared Walsh coefficients, where the weights increase quadratically in the interaction order $k$, corresponding to the fact that higher-order genetic interactions contribute more to local epistasis than lower-order interactions.

Finally, in what follows we will need some basic results on matrices related to $\mathbf{C}$. The first of these is an expression for the Moore-Penrose pseudoinverse of $\mathbf{C}$, $\mathbf{K} = \mathbf{C}^+$, whose entries depend only on $d(i,j)$ and can be calculated individually as

$$\mathbf{K}(i,j) = \sum_{k=2}^{l} \frac{1}{\lambda'_k} w_k(d(i,j)). \tag{8}$$

We also will need a characterization of the ranks of certain submatrices of $\mathbf{C}$ and $\mathbf{K}$. In particular, given an arbitrary subset of sequences $B$ of size $m$ of the set of all possible sequences $\mathcal{S}$, let $I = \mathcal{S} \setminus B$ denote the set of all missing sequences. Without loss of generality, we will order our sequences such that sequences in $B$ come first. Any function on the space of genotypes $\mathbf{f}$ can be written as $\mathbf{f}^T = \begin{bmatrix} \mathbf{f}_B{}^T & \mathbf{f}_I{}^T \end{bmatrix}$. Similarly, we can write $\mathbf{C}$ as

$$\mathbf{C} = \begin{bmatrix} \mathbf{C}_{BB} & \mathbf{C}_{BI} \\ \mathbf{C}_{BI} & \mathbf{C}_{II} \end{bmatrix}. \tag{9}$$

Define a matrix $\mathbf{A} = \begin{bmatrix} \mathbf{I}_m \, \mathbf{0}_{m \times (n-m)} \end{bmatrix}$, so $\mathbf{A}\mathbf{f} = \mathbf{f}_B$. Similarly we have $\mathbf{A}\mathbf{C}\mathbf{A}^T = \mathbf{C}_{BB}$. Next we derive a important result about the rank of the submatrix $\mathbf{C}_{II}$, which will be important for the uniqueness condition proved in Section 2. Note that this result is true for any principal submatrix (a principal submatrix is obtained by deleting the $i$-th rows and $i$-th columns for some values of $i$) of any real-valued positive semi-definite matrix. Here we use $\mathbf{C}$ and $I$ to avoid introducing new notations.

**Lemma 1.** *Let $\mathbf{C}_{\cdot I}$ denote the submatrix of $\mathbf{C}$ containing columns indexed by $I$, then the matrix $\mathbf{C}_{\cdot I}$ and $\mathbf{C}_{II}$ have the same rank for any $I$.*

*Proof.* Let $\mathbf{K} = \mathbf{C}^+$. For any $\mathbf{f} \in \mathrm{col}(\mathbf{C})$, we have $\mathbf{K}\mathbf{C}\mathbf{f} = \mathbf{f}$ and $\mathbf{k}_i^T \mathbf{C}\mathbf{f} = \mathbf{f}(i)$, where $\mathbf{k}_i$ is the $i$-th column of $\mathbf{K}$. Consequently, $\mathbf{C}_{I\cdot}\mathbf{K}\mathbf{C}_{\cdot I} = \mathbf{C}_{II}$. Define a matrix $\mathbf{K}^{\frac{1}{2}}$ by constructing the eigendecomposition of $\mathbf{K}$ and replacing each of its eigenvalues by its square-root. Since the eigenvalues of $\mathbf{K}$ are non-negative, the resulting matrix is also positivesemidefinite and real-valued with the property that $\mathbf{K}^{\frac{1}{2}}\mathbf{K}^{\frac{1}{2}} = \mathbf{K}$. Since

$\mathrm{N}(\mathbf{K}) = \mathrm{N}(\mathbf{K}^{\frac{1}{2}})$ and $\mathrm{col}(\mathbf{C}_{\cdot I}) \cap \mathrm{N}(\mathbf{K}^{\frac{1}{2}}) = \{\mathbf{0}\}$, we have $\mathrm{rank}(\mathbf{K}^{\frac{1}{2}}\mathbf{C}_{\cdot I}) = \mathrm{rank}(\mathbf{C}_{\cdot I})$. Furthermore, since $\mathbf{C}_{I\cdot}\mathbf{K}^{\frac{1}{2}} = (\mathbf{K}^{\frac{1}{2}}\mathbf{C}_{\cdot I})^T$ and $\mathrm{rank}(\mathbf{B}^T\mathbf{B}) = \mathrm{rank}(\mathbf{B})$ for any matrix $\mathbf{B}$, we find

$$\mathrm{rank}(\mathbf{C}_{\cdot I}) = \mathrm{rank}(\mathbf{K}^{\frac{1}{2}}\mathbf{C}_{\cdot I}) = \mathrm{rank}((\mathbf{K}^{\frac{1}{2}}\mathbf{C}_{\cdot I})^T(\mathbf{K}^{\frac{1}{2}}\mathbf{C}_{\cdot I})) = \mathrm{rank}(\mathbf{C}_{I\cdot}\mathbf{K}^{\frac{1}{2}}\mathbf{K}^{\frac{1}{2}}\mathbf{C}_{\cdot I}) = \mathrm{rank}\,\mathbf{C}_{II}. \tag{10}$$

□

### 2 Uniqueness condition for the minimum epistasis interpolation solution

In the main text, we claim that minimum epistasis interpolation has a unique solution if and only if the least squares fit of the corresponding non-epistatic model has a unique solution. Here we prove a stronger result that shows that the epistatic portion of the minimum epistasis interpolation problem is always unique, that any underdetermined component of the solution is limited to the non-epistatic component of the model, that moreover the underdetermined component is exactly shared by the minimum epistasis interpolation and non-epistatic fits. Importantly, whether or not these two problems have a unique solution depends only on the identities of the observed genotypes and does not depend on their actual phenotypic values.

**Proposition 1.** *Let $\widehat{\mathbf{f}}_{add} \in \mathbb{R}^n$ be a non-epistatic fitness landscape fit to the data using ordinary least squares and $\widehat{\mathbf{f}}_{interp} \in \mathbb{R}^n$ a landscape reconstructed using minimum epistasis interpolation. The space of all optimal landscapes for the two methods are given by $\{\widehat{\mathbf{f}}_{add} + \mathbf{X}\mathbf{w} : \mathbf{w} \in \mathrm{N}(\mathbf{X}_B)\}$ and $\{\widehat{\mathbf{f}}_{interp} + \mathbf{X}\mathbf{w} : \mathbf{w} \in \mathrm{N}(\mathbf{X}_B)\}$, respectively. As a consequence, the minimum epistasis interpolation solution is unique if and only if the ordinary least squares problem for the corresponding non-epistatic model has a unique solution.*

*Proof.* Suppose we have observations for some subset $B \subset \mathcal{S}$ of the $n = \alpha^l$ possible genotypes and these observations are given by the vector $\mathbf{y} \in \mathbb{R}^m$. If we fit a non-epistatic model to these data, the model takes the form

$$\mathbf{y} = \mathbf{X}_B\boldsymbol{\beta} + \epsilon, \tag{11}$$

where $\mathbf{X}_B$ is the submatrix of $\mathbf{X}$ containing rows corresponding to $B$. To fit such a model using least squares we must then solve the normal equations

$$(\mathbf{X}_B^T\mathbf{X}_B)\widehat{\boldsymbol{\beta}} = \mathbf{X}_B^T\mathbf{y}. \tag{12}$$

Since $\mathbf{X}_B^T\mathbf{y}$ is in the column space of $\mathbf{X}_B^T\mathbf{X}_B$, this equation always has at least one solution. Let $\widehat{\boldsymbol{\beta}}$ be one such solution, which produces a vector of predicted phenotypes over all of sequence space given by $\widehat{\mathbf{f}}_{\mathrm{add}} = \mathbf{X}\widehat{\boldsymbol{\beta}}$. If $\widehat{\boldsymbol{\beta}}'$ is another solution to Equation 12 then we must have $\widehat{\boldsymbol{\beta}}' - \widehat{\boldsymbol{\beta}} \in \mathrm{N}(\mathbf{X}_B^T\mathbf{X}_B) = \mathrm{N}(\mathbf{X}_B)$ and so the set of all optimal non-epistatic models is given by:

$$\{\widehat{\mathbf{f}}_{\mathrm{add}} + \mathbf{X}\mathbf{w} : \mathbf{w} \in \mathrm{N}(\mathbf{X}_B)\}. \tag{13}$$

Now recall the minimum epistasis interpolation problem

$$\begin{aligned} &\text{minimize } \mathbf{f}^T\mathbf{C}\mathbf{f} \\ &\text{subject to } \mathbf{f}_B = \mathbf{y}. \end{aligned} \tag{14}$$

Expanding $\mathbf{f}^T\mathbf{C}\mathbf{f}$ using Equation 9, we get $\mathbf{f}^T\mathbf{C}\mathbf{f} = \mathbf{f}_B^T\mathbf{C}_{BB}\mathbf{f}_B + 2\mathbf{f}_B^T\mathbf{C}_{BI}\mathbf{f}_I + \mathbf{f}_I^T\mathbf{C}_{II}\mathbf{f}_I$. Setting $\mathbf{f}_B = \mathbf{y}$, we find that the previous equality constrained minimization problem is equivalent to the unconstrained problem in $\mathbf{f}_I$

$$\text{minimize } \frac{1}{2}\mathbf{f}_I^T\mathbf{C}_{II}\mathbf{f}_I + \mathbf{y}^T\mathbf{C}_{BI}\mathbf{f}_I. \tag{15}$$

Since the eigenvalues of $\mathbf{C}$ are nonnegative, all eigenvalues of $\mathbf{C}_{II}$ are bounded below by zero due to the Cauchy interlacing theorem [10], i.e. $\mathbf{C}_{II}$ is positive semi-definite for any $I$. Thus, the function $\frac{1}{2}\mathbf{f}_I^T\mathbf{C}_{II}\mathbf{f}_I + \mathbf{y}^T\mathbf{C}_{BI}\mathbf{f}_I$ is convex. Taking the gradient with respect to $\mathbf{f}_I$ yields $\nabla_{\mathbf{f}_I}(\frac{1}{2}\mathbf{f}_I^T\mathbf{C}_{II}\mathbf{f}_I + \mathbf{y}^T\mathbf{C}_{BI}\mathbf{f}_I) = \mathbf{C}_{II}\mathbf{f}_I + \mathbf{C}_{IB}\mathbf{y}$. Setting this gradient to zero gives us the optimality condition

$$\mathbf{C}_{II}\widehat{\mathbf{f}}_I + \mathbf{C}_{IB}\mathbf{y} = \mathbf{0}. \tag{16}$$

Based on Lemma 1, $\mathrm{col}(\mathbf{C}_{IB})$ is a subspace of $\mathrm{col}(\mathbf{C}_{II})$. Therefore, at least one solution, $\widehat{\mathbf{f}}_I$, exists and $\widehat{\mathbf{f}}_{\text{interp}} = \begin{bmatrix}\mathbf{y}\\ \widehat{\mathbf{f}}_I\end{bmatrix}$ is an optimal landscape reconstructed using the minimum epistasis interpolation. Since if $\widehat{\mathbf{f}}_I'$ is another solution to Equation 16 we must have $\widehat{\mathbf{f}}_I' - \widehat{\mathbf{f}}_I \in \mathrm{N}(\mathbf{C}_{II})$, the space of all optimal landscapes is then given by

$$\{\widehat{\mathbf{f}}_{\text{interp}} + \left[\begin{smallmatrix}\mathbf{0}\\ \mathbf{u}\end{smallmatrix}\right] : \mathbf{u} \in \mathrm{N}(\mathbf{C}_{II})\}. \tag{17}$$

In order to connect this set of solutions to the minimum epistasis interpolation problem to the solutions of the non-epistatic model, we will now show that in fact $\{\left[\begin{smallmatrix}\mathbf{0}\\ \mathbf{u}\end{smallmatrix}\right] : \mathbf{u} \in \mathrm{N}(\mathbf{C}_{II})\} = \{\mathbf{Xw} : \mathbf{w} \in \mathrm{N}(\mathbf{X}_B)\}$. In particular, for any $\mathbf{u} \in \mathrm{N}(\mathbf{C}_{II})$, $\mathbf{C}\left[\begin{smallmatrix}\mathbf{0}\\ \mathbf{u}\end{smallmatrix}\right] = \mathbf{C}_{\cdot I}\mathbf{u} = \left[\begin{smallmatrix}\mathbf{C}_{BI}\mathbf{u}\\ \mathbf{C}_{II}\mathbf{u}\end{smallmatrix}\right] = \mathbf{0}$, since $\mathbf{C}_{II}\mathbf{u} = \mathbf{0}$ and the rows of $\mathbf{C}_{BI}$ are linear combinations of rows of $\mathbf{C}_{II}$ by Lemma 1. So $\left[\begin{smallmatrix}\mathbf{0}\\ \mathbf{u}\end{smallmatrix}\right] \in \mathrm{N}(\mathbf{C}) = \mathrm{col}(\mathbf{X})$ and thus we can find a unique vector $\mathbf{w} \in \mathrm{N}(\mathbf{X}_B)$ such that $\mathbf{Xw} = \left[\begin{smallmatrix}\mathbf{X}_B\\ \mathbf{X}_I\end{smallmatrix}\right]\mathbf{w} = \left[\begin{smallmatrix}\mathbf{0}\\ \mathbf{u}\end{smallmatrix}\right]$. Thus, for any element of the set of solutions given in Equation 17, we can re-express this element in terms of a corresponding $\mathbf{w} \in \mathrm{N}(\mathbf{X}_B)$.

Similarly, we can work in the opposite direction to show that for any $\mathbf{w} \in \mathrm{N}(\mathbf{X}_B)$, we can find a corresponding $\mathbf{u} \in \mathrm{N}(\mathbf{C}_{II})$. In particular, for any $\mathbf{w} \in \mathrm{N}(\mathbf{X}_B)$ we find $\mathbf{Xw} = \left[\begin{smallmatrix}\mathbf{0}\\ \mathbf{X}_I\mathbf{w}\end{smallmatrix}\right]$. Since $\mathbf{Xw} \in \mathrm{N}(\mathbf{C})$, we have $\mathbf{C}(\mathbf{Xw}) = \left[\begin{smallmatrix}\mathbf{C}_{BI}(\mathbf{X}_I\mathbf{w})\\ \mathbf{C}_{II}(\mathbf{X}_I\mathbf{w})\end{smallmatrix}\right] = \mathbf{0}$. As a result $\mathbf{X}_I\mathbf{w} \in \mathrm{N}(\mathbf{C}_{II})$. Therefore, given $\mathbf{w} \in \mathrm{N}(\mathbf{X}_B)$, we have $\mathbf{Xw} = \left[\begin{smallmatrix}\mathbf{0}\\ \mathbf{u}\end{smallmatrix}\right]$ for some $\mathbf{u} \in \mathrm{N}(\mathbf{C}_{II})$.

To summarize, we have shown that the set of optimal minimum epistasis interpolation solutions can be expressed as

$$\{\widehat{\mathbf{f}}_{\text{interp}} + \left[\begin{smallmatrix}\mathbf{0}\\ \mathbf{u}\end{smallmatrix}\right] : \mathbf{u} \in \mathrm{N}(\mathbf{C}_{II})\} = \{\widehat{\mathbf{f}}_{\text{interp}} + \mathbf{Xw} : \mathbf{w} \in \mathrm{N}(\mathbf{X}_B)\}, \tag{18}$$

and likewise that the set of optimal solutions to the non-epistatic model fit by ordinary least squares can be expressed as

$$\{\widehat{\mathbf{f}}_{\text{add}} + \mathbf{Xw} : \mathbf{w} \in \mathrm{N}(\mathbf{X}_B)\} = \{\widehat{\mathbf{f}}_{\text{add}} + \left[\begin{smallmatrix}\mathbf{0}\\ \mathbf{u}\end{smallmatrix}\right] : \mathbf{u} \in \mathrm{N}(\mathbf{C}_{II})\}. \tag{19}$$

It follows immediately that the minimum epistasis interpolation solution is unique if and only if the corresponding non-epistatic model is uniquely determined, or equivalently $\mathbf{X}_B$ has a trivial null space, i.e. $\mathrm{N}(\mathbf{X}_B) = \{\mathbf{0}\}$.

□

### 3 Kernelized solution

Recall the solution to the minimum epistasis interpolation problem provided in the main text:

$$\begin{bmatrix}\widehat{\mathbf{f}}_B\\ \widehat{\mathbf{f}}_I\end{bmatrix} = \begin{bmatrix}\mathbf{y}\\ -(\mathbf{C}_{II})^{-1}\mathbf{C}_{IB}\mathbf{y}\end{bmatrix}. \tag{20}$$

This solution requires solving a linear equation involving the $(n-m)\times(n-m)$ matrix $\mathbf{C}_{II}$, where $n = \alpha^l$ is the size of the genotypic space and $m$ is the number of observations. This approach becomes impractical when $n$ is large and $m$ is much smaller than $n$. However, in this situation, we can still construct an efficient solution by solving a series of linear equations that involve only an $m \times m$ matrix.

In particular, we show below that the solution to the minimum epistasis interpolation problem can always be written in the form

$$\widehat{\mathbf{f}} = \mathbf{K}_{\cdot B}\widehat{\boldsymbol{\alpha}} + \mathbf{X}\widehat{\boldsymbol{\beta}}. \tag{21}$$

for some $\widehat{\boldsymbol{\alpha}} \in \mathbb{R}^m$ and $\widehat{\boldsymbol{\beta}} \in \mathbb{R}^{1+l(\alpha-1)}$ where $\mathbf{K} = \mathbf{C}^+$ is known as the kernel matrix. The existence of a solution of this form is a special case of the semi-parametric representer theorem [11], which is widely used in statistics and machine learning, but for completeness we provide an elementary proof using only linear algebra below.

Given that we can represent $\widehat{\mathbf{f}}$ in this form, instead of solving for $\widehat{\mathbf{f}}$ directly we can find the corresponding $\widehat{\boldsymbol{\alpha}}$ and $\widehat{\boldsymbol{\beta}}$. In fact, we show that $\widehat{\boldsymbol{\alpha}}$ and $\widehat{\boldsymbol{\beta}}$ can be found in the following manner:

1. Construct the matrix $\mathbf{K}_{BB}$ using Equations 4 and 8. Note that this can be done efficiently, since $\mathbf{K}_{BB}(i,j)$ depends only on the Hamming distance $d(i,j)$ so that the entries of $\mathbf{K}_{BB}$ take at most $l+1$ distinct values which can be pre-computed.

2. Solve for the $m$ by $1+l(\alpha-1)$ matrix $\mathbf{Y}$ that satisfies $\mathbf{K}_{BB}\mathbf{Y} = \mathbf{X}_B$ where $\mathbf{Y}$ can be obtained column-wise by solving a system of $m$ equations once for each column of $\mathbf{X}_B$.

3. Solve the equation $\mathbf{Y}^T\mathbf{X}_B\widehat{\boldsymbol{\beta}} = \mathbf{Y}^T\mathbf{y}$ for $\widehat{\boldsymbol{\beta}}$ to obtain the non-epistatic component of the solution.

4. Solve the $m$ equations $\mathbf{K}_{BB}\widehat{\boldsymbol{\alpha}} = \mathbf{y} - \mathbf{X}_B\widehat{\boldsymbol{\beta}}$ for $\widehat{\boldsymbol{\alpha}}$ to obtain the epistatic component of the solution.

To summarize, the kernelized solution involves solving a total of $2 + l(\alpha-1)$ systems of equations with the same $m \times m$ matrix $\mathbf{K}_{BB}$ on the left hand side and one additional system of $1 + l(\alpha-1)$ equations $\mathbf{Y}^T\mathbf{X}_B\widehat{\boldsymbol{\beta}} = \mathbf{Y}^T\mathbf{y}$ that takes a negligible amount of time to solve in the typical regime where $m > 1 + l(\alpha-1)$. In the above method, we have assumed that there is a unique solution $\widehat{\mathbf{f}}$, however as we have shown in Section 2, if the solution is not unique then this is because there are multiple possible non-epistatic components of the solution and we can simply pick one of these to use as $\widehat{\boldsymbol{\beta}}$ to find the unique epistatic component. We have also assumed that there is a unique $\widehat{\boldsymbol{\alpha}}$, i.e. that $\mathbf{K}_{BB}$ is invertible. By Lemma 1, we see that $\mathbf{K}_{BB}$ is invertible if and only if the submatrix of the design matrix for an additive model fit to the unknown genotypes $\mathbf{X}_I$ is of full rank, which will typically be the case in the relevant regime where most genotypes do not have known phenotypes.

We will now derive the above results. Denote col($\mathbf{C}$) the column space of the matrix $\mathbf{C}$, and col$(\mathbf{C})^\perp$ its orthogonal complement with respect to the usual inner product. We find $\mathrm{col}(\mathbf{C})^\perp = \mathrm{N}(\mathbf{C}^T) = \mathrm{N}(\mathbf{C}) = \mathrm{col}(\mathbf{X})$. Therefore, for any $\mathbf{f} \in \mathbb{R}^n$, we can uniquely decompose it as

$$\mathbf{f} = \mathbf{f}^* + \widetilde{\mathbf{f}} = \mathbf{f}^* + \mathbf{X}\boldsymbol{\beta}, \tag{22}$$

where $\mathbf{f}^* \in \mathrm{col}(\mathbf{C})$ is the epistatic component and $\widetilde{\mathbf{f}} = \mathbf{X}\boldsymbol{\beta} \in \mathrm{col}(\mathbf{C})^\perp$ is the non-epistatic component.

Recall the matrix $\mathbf{K}$ defined in Section 1: $\mathbf{K} = \mathbf{C}^+$ so that for $\mathbf{f} \in \mathrm{col}(\mathbf{C})$, $\mathbf{KCf} = \mathbf{f}$ and $\mathbf{k}_i^T\mathbf{Cf} = \mathbf{f}(i)$, where $\mathbf{k}_i$ is the $i$-th column of $\mathbf{K}$. Furthermore, because $\mathbf{K}$ and $\mathbf{C}$ have the same left null space, we have $\mathrm{col}(\mathbf{K}) = \mathrm{col}(\mathbf{C})$. It turns out that for any solution to the minimum epistasis interpolation problem, the epistatic component must in fact lie in a subspace of col($\mathbf{K}$) spanned by the columns of $\mathbf{K}_{\cdot B}$.

**Proposition 2.** *Let $\mathbf{K}_{\cdot B}$ denote the submatrix of $\mathbf{K}$ containing columns corresponding to $B$. Any solution to the minimum epistasis interpolation problem must admit the form*

$$\mathbf{f} = \mathbf{K}_{\cdot B}\boldsymbol{\alpha} + \mathbf{X}\boldsymbol{\beta}. \tag{23}$$

*Proof.* We have already shown that any $\mathbf{f} \in \mathbb{R}^n$ can be decomposed into the form $\mathbf{f} = \mathbf{f}^* + \mathbf{X}\boldsymbol{\beta}$ where $\mathbf{f}^* = \mathbf{f}^* \in \mathrm{col}(\mathbf{C}) = \mathrm{col}(\mathbf{K})$ and so it suffices to show that the epistatic component of any solution can be represented in the form $\mathbf{f}^* = \mathbf{K}_{\cdot B}\boldsymbol{\alpha}$.

To do this, we first perform Gram-Schmidt orthogonalization on the columns of $\mathbf{K}$, $\{\mathbf{k}_1, \cdots, \mathbf{k}_m, \mathbf{k}_{m+1}, \cdots, \mathbf{k}_n\}$, with respect to the inner product defined by $\mathbf{C}$, where without loss of generality we index $\mathbf{K}$ so that the $m$ observed genotypes appear first before the $n-m$ unobserved genotypes. We end up with a new set of vectors, $\{\mathbf{k}'_1, \cdots, \mathbf{k}'_m, \mathbf{k}'_{m+1}, \cdots, \mathbf{k}'_n\}$ that span col($\mathbf{C}$) and are orthogonal in the sense that

$\mathbf{k}_i'^T \mathbf{C}\mathbf{k}_j' = 0$ for $i \neq j$. We can then decompose $\text{col}(\mathbf{C})$ as $\text{col}(\mathbf{C}) = \text{col}(\mathbf{K}) = \text{col}(\mathbf{K}_{\cdot B}) \oplus \text{col}(\mathbf{K}_{\cdot B})^\perp$, where $\text{col}(\mathbf{K}_{\cdot B}) = \text{span}\{\mathbf{k}_1', \cdots, \mathbf{k}_m'\} = \text{span}\{\mathbf{k}_1, \cdots, \mathbf{k}_m\}$, and $\text{col}(\mathbf{K}_{\cdot B})^\perp = \text{span}\{\mathbf{k}_{m+1}', \cdots, \mathbf{k}_n'\}$.

Given this decomposition, any $\mathbf{f}^* \in \text{col}(\mathbf{C})$ can be decomposed as

$$\mathbf{f}^* = \mathbf{K}_{\cdot B}\boldsymbol{\alpha} + \mathbf{v}, \tag{24}$$

with $\mathbf{K}_{\cdot B}\boldsymbol{\alpha} \in \text{col}(\mathbf{K}_{\cdot B})$, $\mathbf{v} \in \text{col}(\mathbf{K}_{\cdot B})^\perp$, so $(\mathbf{K}_{\cdot B}\boldsymbol{\alpha})^T\mathbf{C}\mathbf{v} = 0$. We need to show that if $\mathbf{f}^*$ is the epistatic component of a solution to the minimum epistasis interpolation problem then we must have $\mathbf{v} = \mathbf{0}$.

To see this, suppose to the contrary that $\mathbf{v} \neq \mathbf{0}$. Then consider the alternative epistatic component $\mathbf{f}^{*\prime} = \mathbf{K}_{\cdot B}\boldsymbol{\alpha}$. For any $i \in B$, we find $\mathbf{f}^*(i) = \mathbf{f}^{*T}\mathbf{C}\mathbf{k}_i = (\mathbf{K}_{\cdot B}\boldsymbol{\alpha} + \mathbf{v})^T\mathbf{C}\mathbf{k}_i = (\mathbf{K}_{\cdot B}\boldsymbol{\alpha})^T\mathbf{C}\mathbf{k}_i = \mathbf{f}^{*\prime T}\mathbf{C}\mathbf{k}_i = \mathbf{f}^{*\prime}(i)$, since $\mathbf{v}^T\mathbf{C}\mathbf{k}_i = 0$. Thus $\mathbf{f}^{*\prime}$ and $\mathbf{f}^*$ behave identically on the data. Furthermore, $(\mathbf{f}^* + \mathbf{X}\boldsymbol{\beta})^T\mathbf{C}(\mathbf{f}^* + \mathbf{X}\boldsymbol{\beta}) = (\mathbf{K}_{\cdot B}\boldsymbol{\alpha})^T\mathbf{C}(\mathbf{K}_{\cdot B}\boldsymbol{\alpha}) + \mathbf{v}^T\mathbf{C}\mathbf{v} > (\mathbf{K}_{\cdot B}\boldsymbol{\alpha})^T\mathbf{C}(\mathbf{K}_{\cdot B}\boldsymbol{\alpha}) = (\mathbf{f}^{*\prime} + \mathbf{X}\boldsymbol{\beta})^T\mathbf{C}(\mathbf{f}^{*\prime} + \mathbf{X}\boldsymbol{\beta})$, since $(\mathbf{X}\boldsymbol{\beta})^T\mathbf{C}(\mathbf{X}\boldsymbol{\beta}) = 0$ and we assumed $\mathbf{v} \neq \mathbf{0}$ and $\mathbf{v} \in \text{col}(\mathbf{C})$ so that $\mathbf{v}^T\mathbf{C}\mathbf{v} > 0$. Then $\mathbf{f}^*$ cannot be the epistatic component of the solution to the minimum epistasis interpolation problem because we have constructed an alternative solution $\mathbf{f}^{*\prime}$ with identical behavior on the known genotypes but strictly less epistasis, which is a contradiction. Therefore for the epistatic component of any solution to the minimum epistasis interpolation problem $\mathbf{f}^*$, we must have $\mathbf{v} = \mathbf{0}$.

To summarize, if $\mathbf{f} = \mathbf{K}_{\cdot B}\boldsymbol{\alpha} + \mathbf{v} + \mathbf{X}\boldsymbol{\beta}$ satisfies $\mathbf{f}_B = \mathbf{y}$, then $\mathbf{f}' = \mathbf{K}_{\cdot B}\boldsymbol{\alpha} + \mathbf{X}\boldsymbol{\beta}$ also satisfies this condition but $\mathbf{f}'^T\mathbf{C}\mathbf{f}' \leq \mathbf{f}^T\mathbf{C}\mathbf{f}$. Therefore, the solution to our minimization problem must admit the form $\mathbf{K}_{\cdot B}\boldsymbol{\alpha} + \mathbf{X}\boldsymbol{\beta}$. □

Now that we know that we can express our solution in the form $\widehat{\mathbf{f}} = \mathbf{K}_{\cdot B}\widehat{\boldsymbol{\alpha}} + \mathbf{X}\widehat{\boldsymbol{\beta}}$, we must derive expressions for $\widehat{\boldsymbol{\alpha}}$ and $\widehat{\boldsymbol{\beta}}$. We start by rewriting our minimization problem as

$$\begin{aligned} &\text{minimize } (\mathbf{K}_{\cdot B}\boldsymbol{\alpha} + \mathbf{X}\boldsymbol{\beta})^T\mathbf{C}(\mathbf{K}_{\cdot B}\boldsymbol{\alpha} + \mathbf{X}\boldsymbol{\beta}) \\ &\text{subject to } \mathbf{K}_{BB}\boldsymbol{\alpha} + \mathbf{X}_B\boldsymbol{\beta} = \mathbf{y}, \end{aligned} \tag{25}$$

where we note that $(\mathbf{K}_{\cdot B}\boldsymbol{\alpha} + \mathbf{X}\boldsymbol{\beta})^T\mathbf{C}(\mathbf{K}_{\cdot B}\boldsymbol{\alpha} + \mathbf{X}\boldsymbol{\beta}) = \boldsymbol{\alpha}^T\mathbf{K}_{BB}\boldsymbol{\alpha}$, since $(\mathbf{K}_{\cdot B})^T\mathbf{C}\mathbf{K}_{\cdot B} = \mathbf{K}_{BB}$ and the columns of $\mathbf{X}$ are all in the shared nullspace of $\mathbf{C}$ and $\mathbf{K}$.

Momentarily viewing $\boldsymbol{\beta}$ as fixed, we can take the gradient with respect to $\boldsymbol{\alpha}$ and set this gradient equal to zero, which gives us the condition $\mathbf{K}_{BB}\boldsymbol{\alpha} = \mathbf{y} - \mathbf{X}_B\boldsymbol{\beta}$. Assuming that $\mathbf{K}_{BB}$ is invertible we can thus set $\boldsymbol{\alpha} = (\mathbf{K}_{BB})^{-1}(\mathbf{y} - \mathbf{X}\boldsymbol{\beta})$ and convert our problem into the unconstrained minimization problem:

$$\min_{\boldsymbol{\beta} \in \mathbb{R}^{1+l(\alpha-1)}} (\mathbf{y} - \mathbf{X}_B\boldsymbol{\beta})^T(\mathbf{K}_{BB})^{-1}(\mathbf{y} - \mathbf{X}_B\boldsymbol{\beta}). \tag{26}$$

This problem is strictly convex in $\mathbf{X}_B\boldsymbol{\beta}$ since $(\mathbf{K}_{BB})^{-1}$ has all positive eigenvalues again due to the Cauchy interlacing theorem. If $\mathbf{X}_B$ has full column rank, then it is strictly convex in $\boldsymbol{\beta}$. Taking the gradient with respect to $\boldsymbol{\beta}$ and setting this gradient to zero, we arrive at the optimality condition

$$(\mathbf{X}_B)^T(\mathbf{K}_{BB})^{-1}\mathbf{X}_B\widehat{\boldsymbol{\beta}} = (\mathbf{X}_B)^T(\mathbf{K}_{BB})^{-1}\mathbf{y}, \tag{27}$$

where the solution $\widehat{\boldsymbol{\beta}}$ is unique since the matrix $(\mathbf{X}_B)^T(\mathbf{K}_{BB})^{-1}\mathbf{X}_B$ is positive definite based on our assumption that $\mathbf{X}_B$ has full column rank. The corresponding optimal $\widehat{\boldsymbol{\alpha}}$ is then given by

$$\widehat{\boldsymbol{\alpha}} = (\mathbf{K}_{BB})^{-1}(\mathbf{y} - \mathbf{X}_B\widehat{\boldsymbol{\beta}}). \tag{28}$$

As described above, in practice we generally do not directly calculate $(\mathbf{K}_{BB})^{-1}$. Instead, we solve $1 + l(\alpha - 1)$ equations involving $\mathbf{K}_{BB}$ to find a matrix $\mathbf{Y}$ so that $\mathbf{X}_B = \mathbf{K}_{BB}\mathbf{Y}$. We can then solve Equation 27 as $(\mathbf{X}_B)^T\mathbf{Y}\widehat{\boldsymbol{\beta}} = \mathbf{Y}^T\mathbf{y}$ for $\widehat{\boldsymbol{\beta}}$ and then solve one more set of equation involving $\mathbf{K}_{BB}$ (i.e. $\mathbf{K}_{BB}\widehat{\boldsymbol{\alpha}} = \mathbf{y} - \mathbf{X}_B\widehat{\boldsymbol{\beta}}$) to find $\widehat{\boldsymbol{\alpha}}$.

### 4 Limiting solutions of a general minimization problem

In the main text, we state that the classical non-epistatic model fit by least squares and the minimum epistasis interpolation models can be viewed as two ends of a continuum of models that are parametrized by the relative importance given to minimizing the mean squared epistatic coefficient as compared to the mean squared error. Here we provide a characterization of this set of models. Besides establishing that minimum epistasis interpolation and the non-epistatic model are two limiting cases of this broader class of models, in passing we also derive several other characteristics of this broader class of models.

The most important of these characteristics is that given the fit of the model on the observed genotypes (i.e. $\widehat{\mathbf{f}}_B$), the out-of-sample predictions of the model are equivalent to doing minimum epistasis interpolation based on these fit values. As a result, the solutions within this broader class share the geometric properties of the minimum epistasis interpolation solution described in the main text, such as their relation to the average non-epistatic solution, average predictions at distances 1 and 2, etc. We also establish that the uniqueness conditions and character of the underdetermination for this broader class of models is the same as that described in Section 2 of this SI document, i.e. any underdetermination is limited to the non-epistatic component of the solution.

We now describe this general minimization problem. In particular, for $0 < p < 1$ consider the minimization problem

$$\text{minimize} \quad p\mathbf{f}^T\mathbf{C}\mathbf{f} + (1-p)\|\mathbf{A}\mathbf{f} - \mathbf{y}\|^2. \tag{29}$$

Note that this general formulation differs from our minimum epistasis problem 14 mainly in the second term, where in 14 we restrict the solution to be in the subset of function that match the data exactly, whereas here we allow deviation from the data and the solution results from the balance between the smoothness term $\mathbf{f}^T\mathbf{C}\mathbf{f}$ and the sum of squared difference $\|\mathbf{A}\mathbf{f} - \mathbf{y}\|^2$ with their relative importance parameterized by $p$. Note that both terms are quadratic in $\mathbf{f}$ and non-negative, so we know immediately that at least one solution exists. We now turn to demonstrating that the limiting solutions as $p \to 0$ and $p \to 1$ coincide with the minimum epistasis solution and the non-epistatic model fit by ordinary least squares solution, respectively.

First consider the case where we have observed phenotypes for all sequences, that is $B = \mathcal{S}$. Taking the gradient with respect to $\mathbf{f}$ in Equation 29 and setting the result to zero, we find the minimizer as the solution to

$$\left(\frac{p}{1-p}\mathbf{C} + \mathbf{I}\right)\widehat{\mathbf{f}} = \mathbf{y}. \tag{30}$$

Since $\mathbf{C} = \sum_{k=2}^{l} \frac{\alpha^2}{2s}(k^2-k)\mathbf{W}_k$ and $\mathbf{I} = \sum_{k=0}^{l} \mathbf{W}_k$, we have

$$\frac{p}{1-p}\mathbf{C} + \mathbf{I} = \mathbf{W}_0 + \mathbf{W}_1 + \sum_{k=2}^{l}(\frac{p}{1-p}\frac{\alpha^2}{2s}(k^2-k)+1)\mathbf{W}_k. \tag{31}$$

This matrix has all nonzero eigenvalues for $0 < p < 1$, therefore has an inverse given by

$$(\frac{p}{1-p}\mathbf{C} + \mathbf{I})^{-1} = \mathbf{W}_0 + \mathbf{W}_1 + \sum_{k=2}^{l}(\frac{p}{1-p}\frac{\alpha^2}{2s}(k^2-k)+1)^{-1}\mathbf{W}_k \tag{32}$$

since $\mathbf{W}_k = \sum_{i=1}^{m_k} \mathbf{u}_{k_i}(\mathbf{u}_{k_i})^T$, where the $\mathbf{u}_{k_i}$'s are orthonormal eigenvectors for $V_k$ and $\mathbf{W}_0 + \mathbf{W}_1$ is the projection matrix for the subspace $V_0 \oplus V_1$. Moreover, because the columns of the design matrix $\mathbf{X}$ also spans the subspace $V_0 \oplus V_1$ and are linearly independent, we also have $\mathbf{W}_0 + \mathbf{W}_1 = \mathbf{X}(\mathbf{X}^T\mathbf{X})^{-1}\mathbf{X}^T$.

Together Equation 30 and the inverse provided in Equation 32 constitute a formal solution for any $p$ and thus we turn to characterizing the two limiting cases. Considering first the case where $p \to 0$, we find $\lim_{p\to 0}(\frac{p}{1-p}\frac{\alpha^2}{2s}(k^2-k)+1)^{-1} = 1$. Therefore, we have $(\frac{p}{1-p}\mathbf{C}+\mathbf{I})^{-1} \to \mathbf{I}$ and $\widehat{\mathbf{f}} \to \mathbf{y}$ as $p \to 0$, which is the same as the (in this case, trivial) solution to the minimum epistasis interpolation problem. For the case where $p \to 1$, we have for $k \geq 2$, $\lim_{p\to 1}(\frac{p}{1-p}\frac{\alpha^2}{2s}(k^2-k)+1)^{-1} = 0$. Thus $\widehat{\mathbf{f}} \to \mathbf{X}(\mathbf{X}^T\mathbf{X})^{-1}\mathbf{X}^T\mathbf{y}$ as $p \to 1$, which is identical to the solution of the non-epistatic model fit by least squares.

Having described the two limiting solutions for the special case where phenotypic data for all genotypes is available, we now consider the case where we have observations for a strict subset of all possible sequences. Rearranging Equation 29 gives

$$\text{minimize} \quad \mathbf{f}^T \left( p\mathbf{C} + (1-p) \begin{bmatrix} \mathbf{I}_m & \mathbf{0} \\ \mathbf{0} & \mathbf{0} \end{bmatrix} \right) \mathbf{f} - (1-p)2\mathbf{y}^T\mathbf{A}\mathbf{f} + (1-p)\|\mathbf{y}\|^2, \tag{33}$$

and taking the gradient with respect to $\mathbf{f}$ and setting it equal to zero we find the optimality condition:

$$(p \begin{bmatrix} \mathbf{C}_{BB} & \mathbf{C}_{BI} \\ \mathbf{C}_{IB} & \mathbf{C}_{II} \end{bmatrix} + (1-p) \begin{bmatrix} \mathbf{I}_m & \mathbf{0} \\ \mathbf{0} & \mathbf{0} \end{bmatrix}) \begin{bmatrix} \widehat{\mathbf{f}}_B \\ \widehat{\mathbf{f}}_I \end{bmatrix} = (1-p) \begin{bmatrix} \mathbf{y} \\ \mathbf{0} \end{bmatrix}. \tag{34}$$

Let $\widehat{\mathbf{f}}$ be a solution to this equation, then the set of all solutions is given by $S = \{\widehat{\mathbf{f}} + \mathbf{v} : \mathbf{v} \in \mathrm{N}(p\mathbf{C} + (1-p)\left[\begin{smallmatrix} \mathbf{I}_m & \mathbf{0} \\ \mathbf{0} & \mathbf{0} \end{smallmatrix}\right])\} = \{\widehat{\mathbf{f}} + \mathbf{v} : \mathbf{v} \in \mathrm{N}(\mathbf{C}) \cap \mathrm{N}(\left[\begin{smallmatrix} \mathbf{I}_m & \mathbf{0} \\ \mathbf{0} & \mathbf{0} \end{smallmatrix}\right])\}$. Since a vector in $\mathrm{N}(\mathbf{C})$ must take the form $\mathbf{X}\mathbf{w}$ for some $\mathbf{w}$, and a vector in $\mathrm{N}(\left[\begin{smallmatrix} \mathbf{I}_m & \mathbf{0} \\ \mathbf{0} & \mathbf{0} \end{smallmatrix}\right])$ must take the form $\left[\begin{smallmatrix} \mathbf{0} \\ \mathbf{z} \end{smallmatrix}\right]$, for some vector $\mathbf{z} \in \mathbb{R}^{n-m}$, we find $\mathbf{v} \in \mathrm{N}(\mathbf{C}) \cap \mathrm{N}(\left[\begin{smallmatrix} \mathbf{I}_m & \mathbf{0} \\ \mathbf{0} & \mathbf{0} \end{smallmatrix}\right])$ must admit the form $\mathbf{v} = \mathbf{X}\mathbf{w}$ for $\mathbf{w} \in \mathrm{N}(\mathbf{X}_B)$. Consequently, we have $S = \{\widehat{\mathbf{f}} + \mathbf{X}\mathbf{w} : \mathbf{w} \in \mathrm{N}(\mathbf{X}_B)\}$ and thus the solution is unique precisely when the minimum epistasis interpolation and non-epistatic models have unique solutions, i.e. when $\mathrm{N}(\mathbf{X}_B) = \{\mathbf{0}\}$, and any undertermination is independent of $p$ and only influences the non-epistatic component of the landscape in a manner completely parallel with the results of Section 2.

Now rewrite Equation 34 as two equations

$$\begin{cases} p\mathbf{C}_{BB}\widehat{\mathbf{f}}_B + p\mathbf{C}_{BI}\widehat{\mathbf{f}}_I + (1-p)\widehat{\mathbf{f}}_B = (1-p)\mathbf{y} \\ \mathbf{C}_{IB}\widehat{\mathbf{f}}_B + \mathbf{C}_{II}\widehat{\mathbf{f}}_I = \mathbf{0}. \end{cases} \tag{35}$$

The second equation tells us that given $\widehat{\mathbf{f}}_B$, the set of optimal $\widehat{\mathbf{f}}_I$ is specified by the solutions to $\mathbf{C}_{II}\widehat{\mathbf{f}}_I = -\mathbf{C}_{IB}\widehat{\mathbf{f}}_B$, which is identical to the solution of the minimum epistasis interpolation problem with observations $\widehat{\mathbf{f}}_B$ instead of $y$. Thus, given the in-sample predictions $\widehat{\mathbf{f}}_B$, the out-of-sample predictions for any $p$ are given by the minimum epistasis interpolation extension of the in-sample predictions and hence satisfy the geometric characteristics described in the main text.

Having characterized the nature of the solutions for this problem, we now characterize the behavior of the solution in the limits of $p \to 0$ and $p \to 1$.

To derive the limit for $p \to 0$, we begin by solving the second system of equations in Equation 35. Recall that this system of equations always has at least one solution since $\mathrm{col}(\mathbf{C}_{IB}) \subset \mathrm{col}(\mathbf{C}_{II})$. Furthermore, denoting $\mathbf{C}_{II}^+$ the pseudoinverse of $\mathbf{C}_{II}$, we can verify $\widehat{\mathbf{f}}_I = -(\mathbf{C}_{II})^+\mathbf{C}_{IB}\widehat{\mathbf{f}}_B$ is a solution, so that the set of all optimal $\widehat{\mathbf{f}}_I$ is given by $\{-(\mathbf{C}_{II})^+\mathbf{C}_{IB}\widehat{\mathbf{f}}_B + \mathbf{u} : \mathbf{u} \in \mathrm{N}(\mathbf{C}_{II})\}$. Now, bring any vector in $\{-(\mathbf{C}_{II})^+\mathbf{C}_{IB}\widehat{\mathbf{f}}_B + \mathbf{u} : \mathbf{u} \in \mathrm{N}(\mathbf{C}_{II})\}$ into the first equation of Equation 35, we get

$$(p\mathbf{H} + \mathbf{I}_m)\widehat{\mathbf{f}}_B = (1-p)\mathbf{y}, \tag{36}$$

where we set $\mathbf{H} = \mathbf{C}_{BB} - \mathbf{C}_{BI}(\mathbf{C}_{II})^+\mathbf{C}_{IB} - \mathbf{I}_m$ and note that $\mathbf{u}$ disappears from this equation since $\mathbf{u} \in \mathrm{N}(\mathbf{C}_{II}) \supset \mathrm{N}(\mathbf{C}_{BI})$.

For $p$ approximating 0, we can write $\widehat{\mathbf{f}}_B = (p\mathbf{H} + \mathbf{I}_m)^{-1}(1-p)\mathbf{y}$, since $(p\mathbf{H} + \mathbf{I}_m)^{-1}$ always exits for sufficiently small $p$ due to the Gershgorin circle theorem [10]. Since $p\mathbf{H} + \mathbf{I}_m \to \mathbf{I}_m$ as $p \to 0$ and matrix inversion is a continuous function over the set of invertible matrices, we find $(p\mathbf{H} + \mathbf{I}_m)^{-1} \to \mathbf{I}_m$. Consequently, $\lim_{p\to 0}\widehat{\mathbf{f}}_B = \lim_{p\to 0}(p\mathbf{H} + \mathbf{I}_m)^{-1}(1-p)\mathbf{y} = \mathbf{y}$. Correspondingly, the set of all optimal

$\widehat{\mathbf{f}}_I$ is given by solutions to the equation $\mathbf{C}_{IB}\mathbf{y} + \mathbf{C}_{II}\widehat{\mathbf{f}}_I = \mathbf{0}$, which is exactly the minimum epistasis interpolation solution (see Equation 20).

To derive the limiting solution for $p \to 1$, we note that the result in Proposition 2 also applies to Equation 29 using a similar argument, i.e. the solution must admit the form $\mathbf{f} = \mathbf{K}_{\cdot B}\boldsymbol{\alpha} + \mathbf{X}\boldsymbol{\beta}$ because any candidate solution $\mathbf{f}$ not of this form can be modified to have a smaller value for $\mathbf{f}^T\mathbf{C}\mathbf{f}$ without increasing $\|\mathbf{A}\mathbf{f} - \mathbf{y}\|^2$. Notes also, that if the columns of $\mathbf{K}_{\cdot B}$ are not linearly independent, we can choose a subset of linearly independent columns corresponding to a subset $B' \subset B$, so that $\operatorname{rank}\mathbf{K}_{\cdot B} = \operatorname{rank}\mathbf{K}_{\cdot B'}$ to ensure $\mathbf{K}_{B'B'}$ is invertible using Proposition 1.

Substituting $\mathbf{f}$ in Problem 29 with this alternative form, taking the gradient with respect to $\widehat{\boldsymbol{\alpha}}$ and $\widehat{\boldsymbol{\beta}}$ and setting the results equal to zero, we have again have two sets of linear equations as the optimality condition:

$$\begin{cases} p\mathbf{K}_{BB}\widehat{\boldsymbol{\alpha}} + (1-p)\mathbf{K}_{BB}\mathbf{K}_{BB}\widehat{\boldsymbol{\alpha}} + (1-p)\mathbf{K}_{BB}\mathbf{X}_B\widehat{\boldsymbol{\beta}} - (1-p)\mathbf{K}_{BB}\mathbf{y} = 0 \\ (\mathbf{X}_B)^T\mathbf{X}_B\widehat{\boldsymbol{\beta}} + \mathbf{X}_B^T\mathbf{K}_{BB}\widehat{\boldsymbol{\alpha}} - \mathbf{X}_B^T\mathbf{y} = 0. \end{cases} \tag{37}$$

For fixed $\widehat{\boldsymbol{\alpha}}$, the set of optimal $\widehat{\boldsymbol{\beta}}$ is given by the solution to the second equation, $\{((\mathbf{X}_B)^T\mathbf{X}_B)^+\mathbf{X}_B^T(\mathbf{y} - \mathbf{K}_{BB}\widehat{\boldsymbol{\alpha}}) + \mathbf{w} : \mathbf{w} \in \mathrm{N}(\mathbf{X}_B)\}$. (We note more generally that this result means that for any value of $0 < p < 1$, the non-epistatic component of the model is equal to a non-epistatic model fit by least squares to the residuals of the epistatic component). Now, bringing any element in this set into the first equation in Equation 37, we get

$$(p\mathbf{I}_m + (1-p)(\mathbf{I}_m - \mathbf{P})\mathbf{K}_{BB})\widehat{\boldsymbol{\alpha}} = (1-p)(\mathbf{I}_m - \mathbf{P})\mathbf{y}, \tag{38}$$

where we have set $\mathbf{P} = \mathbf{X}_B((\mathbf{X}_B)^T\mathbf{X}_B)^+\mathbf{X}_B^T$. Since $\lim_{p\to 1}(1-p)(\mathbf{I}_m - \mathbf{P})\mathbf{y} = \mathbf{0}$ and $\lim_{p\to 1}(p\mathbf{I}_m + (1-p)(\mathbf{I}_m - \mathbf{P})\mathbf{K}_{BB}) = \mathbf{I}_m$, by the continuity of matrix inversion we find $\lim_{p\to 1}\widehat{\boldsymbol{\alpha}} = \mathbf{0}$. Bringing this solution into the second equation in Equation 37, we see that the set of all optimal $\widehat{\boldsymbol{\beta}}$ is given by the solutions to $(\mathbf{X}_B)^T\mathbf{X}_B\widehat{\boldsymbol{\beta}} = \mathbf{X}_B^T\mathbf{y}$, which is simply the normal equations for the non-epistatic model under ordinary least squares (see Equation 12). Thus, we have shown that the limiting solutions as $p \to 0$ and $p \to 1$ to this generalized minimization problem are given by the minimum epistasis interpolation solution and the classical non-epistatic model, respectively.

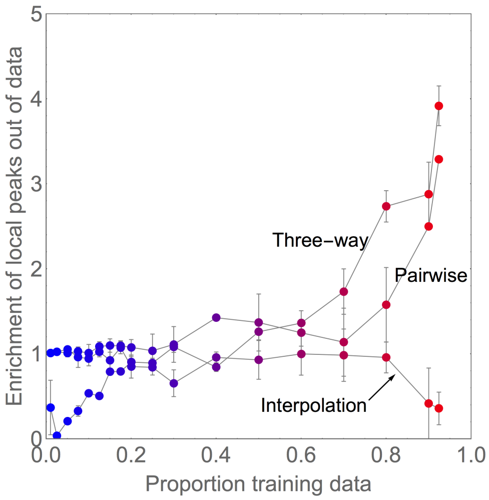


Supplemental Figure 1: Enrichment of local peaks out of data for models fit to the GB1 combinatorial mutagenesis dataset. Pairwise and three-way interaction models were fit using regularized regression with regularization parameters chosen by 10-fold cross-validation. Enrichment score is calculated as (# of local peaks out of data/# of sequences out of data)/(# of local peaks in the data/# of sequences in the data); error bars indicate standard error.

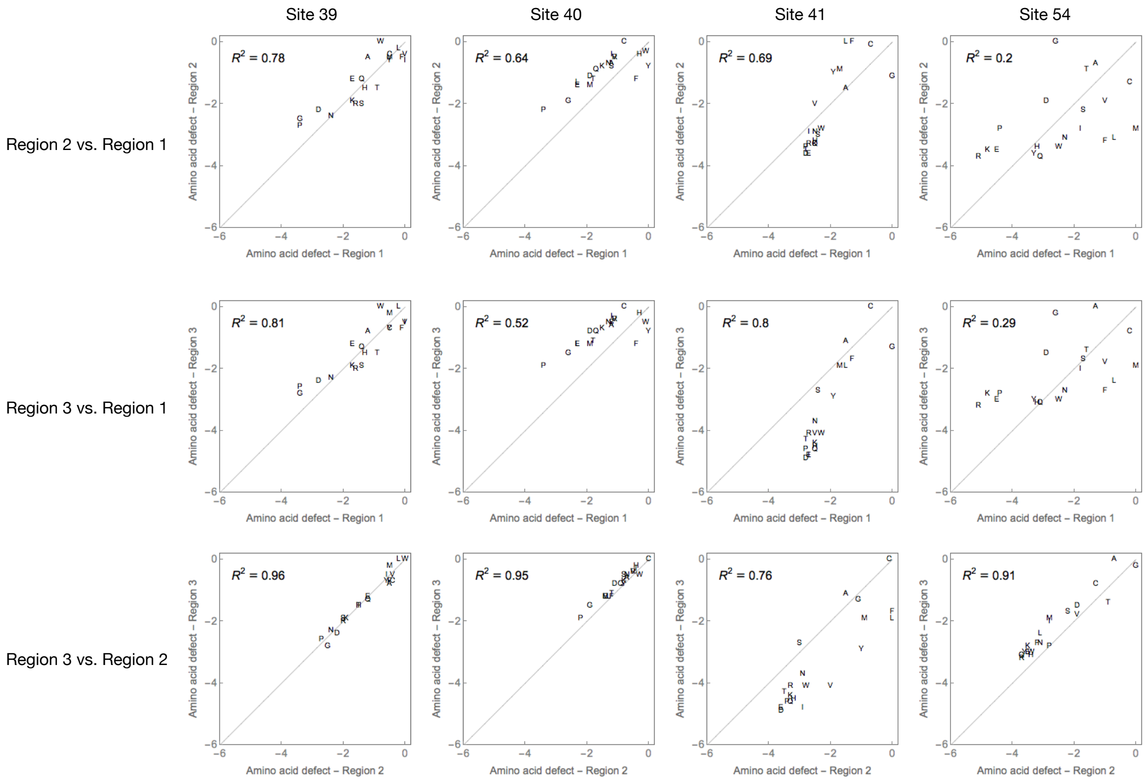


Supplementary Figure 2: Comparison of mean amino acid defects between the three high fitness regions in the GB1 visualization. Letters represent mutations in amino acids. For each site in each genotype in each region, we calculate the effect of replacing the resident amino acid by each of the 20 possible amino acids, average these mutational effects over all genotypes in the region, and subtract the maximum effect for each site to calculate the mean region-specific fitness defect conferred by each amino acid at each site. Deviations from the line $y = x$ indicate differences in amino acid preference between regions.
